# Vitamin D Sufficiency Enhances Differentiation of Patient-Derived Prostate Epithelial Organoids

**DOI:** 10.1101/2020.07.17.208694

**Authors:** Tara McCray, Julian Pacheco, Bethany Baumann, Michael J Schlicht, Klara Valyi-Nagy, Larisa Nonn

## Abstract

Vitamin D is an essential steroid hormone that regulates systemic calcium homeostasis and cell fate decisions, including differentiation in many cell types. The prostate gland is hormonally regulated, requiring steroids for proliferation and terminal differentiation of secretory luminal cells. Vitamin D deficiency is associated with higher risk of lethal prostate cancer with an aggressive dedifferentiated pathology, linking vitamin D sufficiency to epithelial differentiation homeostasis. To determine regulation of prostate epithelial differentiation by vitamin D status, patient-derived benign prostate epithelial organoids were maintained in vitamin D deficient (vehicle) or sufficient (10 nM 1,25-dihydroxyvitamin D) conditions and assessed by phenotype and single cell RNA sequencing. Mechanistic validation demonstrated that vitamin D sufficiency promoted organoid growth and accelerated differentiation of lineage-committed cells by inhibiting canonical Wnt activity and Wnt family member DKK3. Wnt dysregulation is a known contributor to aggressive prostate cancer, thus these findings further link vitamin D deficiency to lethal disease.

## INTRODUCTION

Vitamin D is a misnomer as it is not a vitamin, but rather a vital steroid, synthesized in the skin following UV exposure. Circulating pro-hormone is further metabolized to the active hormone calcitriol [1,25-dihydroxyvitamin D (1,25D)] which controls systemic calcium homeostasis and locally regulates cell fate decisions. In target cells, 1,25D binds to the vitamin D receptor (VDR), a classical steroid hormone receptor that associates with vitamin D response elements on DNA to control gene expression (Feldman et al., 2014). Vitamin D regulates at least 3% of the genome (Bouillon et al., 2008) and ChIP sequencing for VDR-bound-DNA in prostate epithelial cells revealed binding at more than 3,000 protein coding genes, and over 1,000 noncoding sites (Baumann et al., 2019; Fleet et al., 2019). Genes regulated by vitamin D are involved in proliferation, differentiation, and apoptosis pathways (Feldman et al., 2014). Vitamin D metabolites are pro-differentiating in a variety of cell and tissue types, including keratinocytes (MacLaughlin et al., 1990), intestinal villi (Peregrina et al., 2015; Spielvogel et al., 1972), cardiomyocytes (Hlaing et al., 2014), odontoblasts (Mucuk et al., 2017), placenta (Hutabarat et al., 2018), and macrophages (Abe et al., 1981; James et al., 1997). While some of these studies explore 1,25D action in benign cells, the majority of reports focus on vitamin D inhibition of cancer cell growth and tumor progression (Aguilera et al., 2007; Banks and Holick, 2015; Holick et al., 2007; Larriba et al., 2011; Tavera-Mendoza et al., 2017; Yang et al., 2017), and very little is known about the pro-differentiating activity of vitamin D in benign prostate.

Other hormones are essential for prostate differentiation; retinoic acid regulates prostate bud formation during development (Bryant et al., 2014) and androgen regulates terminal differentiation of luminal epithelial cells (Prins and Lindgren, 2015; Wright et al., 1996). The prostate has robust expression of VDR and the enzymes required for local production of 1,25D from the circulating pro-hormone, 25-hydroxyvitamin D (Peehl et al., 2004), indicating that the hormone vitamin D is active in the gland as well. The benign prostate of castrated rats supplemented with androgen and 1,25D showed increased number of secretory vesicles in regenerated luminal cells compared to androgen alone (Konety et al., 1996), supporting improved differentiation with vitamin D. In prostate cancer cell lines, treatment with a vitamin D analog increased E-cadherin expression (Campbell et al., 1997). While these two studies suggest that vitamin D enhanced prostatic differentiation by expression of a more “normal” phenotype, the mechanism was not determined.

Hormone dysregulation, such as androgen and androgen receptor, contributes to prostate cancer (PCa) initiation and progression (Karantanos et al., 2013). Similar observations have been made for vitamin D and VDR; well-differentiated prostate tumors have high VDR expression, whereas high Gleason grade, poorly-differentiated tumors have low VDR expression (Hendrickson et al., 2011). The relationship between vitamin D dysregulation and aggressive disease is supported by rodent models, where prostatic VDR deletion within the TgAPT mouse model of prostate carcinogenesis increased adenocarcinoma foci number and area (Fleet et al., 2019). The relationship is also observed in patients, where vitamin D deficiency is associated with risk of aggressive PCa (Fang et al., 2011; Giovannucci et al., 2006; Murphy et al., 2014; Ramakrishnan et al., 2019; Studzinski and Moore, 1995). Vitamin D deficiency is especially pertinent to PCa patients who are frequently deficient due to high skin melanin content, such as African Americans, (Murphy et al., 2014) or lack of sun exposure (Gilbert et al., 2009), such as the elderly (Elshazly et al., 2017). This observation was originally identified as distance from equator increasing the risk of prostate cancer mortality (Hanchette and Schwartz, 1992; Studzinski and Moore, 1995), and diagnosis during summer improving PCa prognosis (Robsahm et al., 2004). Clinical trials using vitamin D supplementation in patients with existing PCa have had both promising (Marshall et al., 2012; Woo et al., 2005) and null results (Chandler et al., 2014). However, patients with PCa are known to have low expression of VDR (Hendrickson et al., 2011) supporting that vitamin D supplementation is more useful as a chemopreventive agent rather than a PCa treatment. Recently, the VITAL study assessed 2000 IU/day vitamin D supplementation in healthy subjects to explore cancer incidence and found a decreased risk of cancer in the normal BMI group (Manson et al., 2019).

Given that the reliance of the prostate on hormone signaling for differentiation, the high activity of the hormone vitamin D in the gland, and the association of vitamin D deficiency with risk of aggressive PCa (Murphy et al., 2014), we hypothesized that vitamin D promotes prostatic epithelial differentiation and its deficiency contributes to the loss of epithelial homeostasis observed in aggressive PCa. To explore the effects of deficient and sufficient 1,25D on prostate epithelium, patient-derived organoids were used to model epithelial differentiation (Drost et al., 2016) in physiologically relevant levels of the hormone. We previously optimized culture conditions for clonal growth of organoids from single cells (Richards et al., 2019) and characterized prostate organoids using single-cell RNA sequencing (scRNAseq) (McCray et al., 2019a).

## RESULTS

### Sufficient 1,25D supports growth of benign patient-derived prostate epithelial organoids

To determine the effect of 1,25D on organoid phenotype, we grew organoids derived from a benign region of radical prostatectomy tissue from a single patient in media supplemented with 10 nM 1,25D, 50 nM 1,25D (vitamin D-sufficient) or vehicle control (vitamin D-deficient, <0.01% EtOH). VDR was found to be expressed in all patient-derived PrE cells examined (**Supplemental Figure S1A**), including patient AA2. The clonal origin of organoids within this model has been previously demonstrated by our group through time-lapse microscopy studies and lineage tracing with fluorescent proteins (Richards et al., 2019). Similar to other reports (Barros-Silva et al., 2018; Wang et al., 2015), we observed organoids with heterogeneous morphologies that include solid spheres and acinar/tubule-like structures, with a small percentage of translucent spheres (**Figure 1A**). Both concentrations of 1,25D produced visibly larger organoid areas compared to the vitamin D-deficient control organoids (**Figure 1A**). The 10 nM dosage was selected for subsequent studies, as it strongly upregulated the VDR-response gene *CYP24A1* (Chen and DeLuca, 1995; Zierold et al., 1995) (**Supplemental Figure S1B**) and is a biologically relevant concentration; equivalent to ∼4 ng/mL vitamin D (circulating levels of active and inactive forms in serum are 50 pg/mL and 40 ng/mL, respectively) (Richards et al., 2017). Studies were expanded to include a larger and more diverse group of 10 patient-derived PrE cells (**Table 1**). Vitamin D sufficiency increased the percentage of translucent and acinar organoids, which are considered more differentiated morphologies (**Figure 1B**). Organoid size was significantly increased by 1,25D (**Figure 1C & D**, **Table 1**). The magnitude of the 1,25D-induced size increase varied between patients, highlighting the model’s preservation of patient heterogeneity (**Figure 1 C & D**). Flow cytometry of day 14 organoids for luminal cell marker CD26 (Henry et al., 2018) and basal-progenitor colony-forming cell marker CD49f (Yamamoto et al., 2012) revealed a noticeable shift in CD49f cells at day 14 with 1,25D treatment (blue) as compared to control (red) (**Figure 1E**). Overall, 1,25D significantly increased organoid size and promoted differentiation away from a CD49f basal-progenitor phenotype, indicating improved growth with vitamin D.

**Figure 1.**
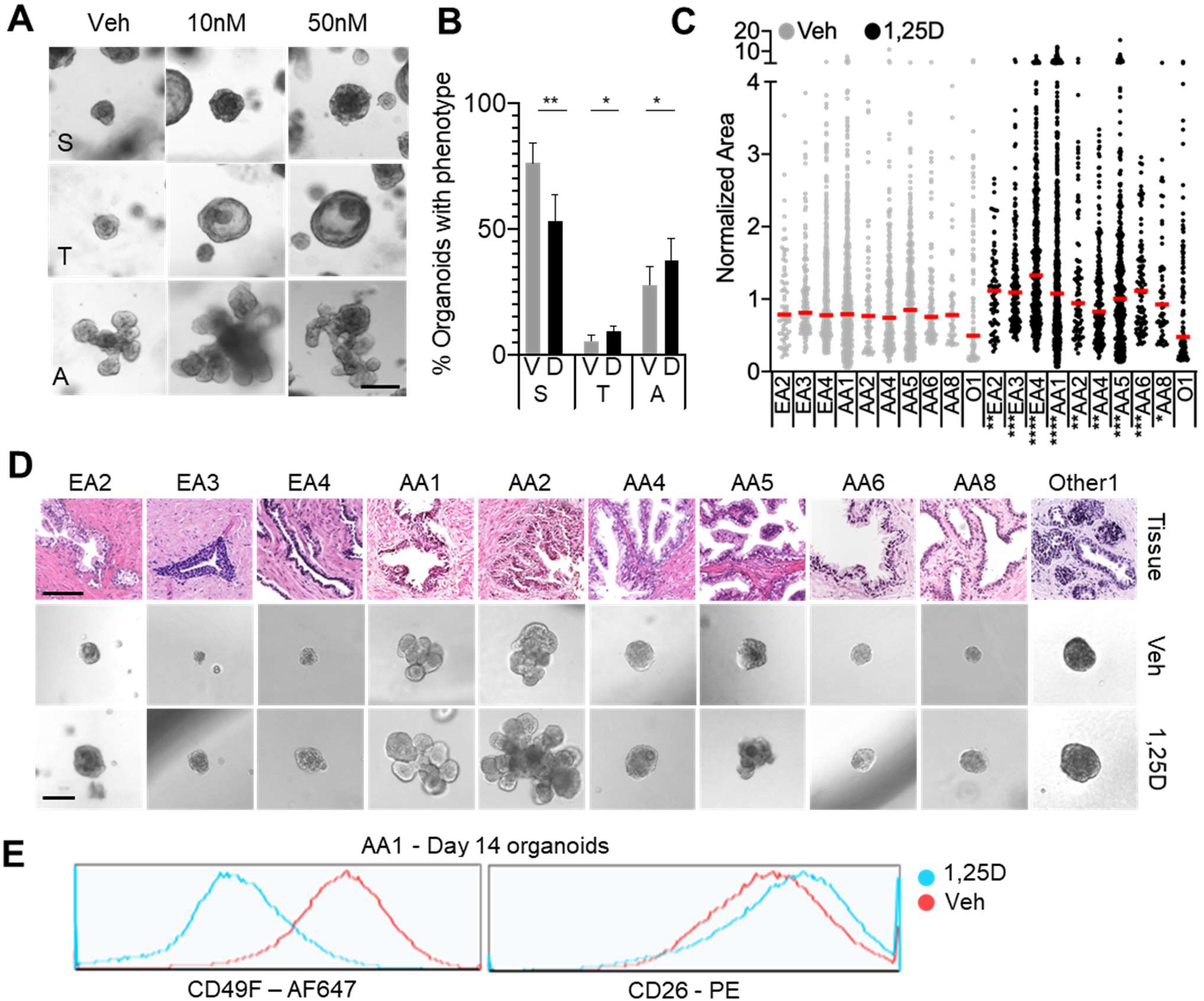
**Sufficient 1,25D supports growth of benign patient-derived prostate epithelial organoids** (see also Figure S1). (A) AA2 organoids grown in vehicle (Veh, <0.01% EtOH), 10 nM or 50 nM 1,25D until day 23. Three representative images are shown for each condition to illustrate heterogeneity of organoid morphology: solid (S), translucent (T), and acinar (A) structures were observed. Scale bar = 200 μm. (B) Percentage of organoids with phenotype after culture in deficient 1,25D (V, vehicle), or sufficient 1,25D (D, 10 nM 1,25D). Solid (S), translucent (T), and acinar (A) organoids were counted per well in n ≥ 3 wells for 10 patients and the percentage of each were calculated. Organoids derived from 7 patients were capable of forming all three types of morphologies, only those percentages were averaged and reported here, error bars represent the standard error of the mean. (*p < 0.1; **p < 0.05) (C) Relative area of PrE organoids grown in vehicle (Veh) or 10 nM 1,25D (1,25D) grown for approximately 14 days, showing ≥ 3 replicate wells per patient for each condition, repeated 1-4 times per patient as listed on **Table 1**. Area is normalized to the mean of the vehicle per patient per experiment, each dot represents an organoid. Patients represent biological replicates. (AA, African American; EA, European American; O, Other, not African or European descent, ancestries were patient self-declared; *p < 0.1; **p < 0.05; ***p < 0.01, ****p < 0.001). (D) Representative images of day-14 organoids grown in deficient (Veh, middle) or sufficient vitamin D (1,25D, lower) and corresponding H&E staining for patient tissue from which organoids were derived (upper), scale bar = 200 μm for organoids and 100 μm for tissue. (E) Overlay of flow histograms for luminal marker CD26 (right) and basal-progenitor marker CD49f (left) in AA1 organoids grown until day 14 in vehicle (red, Veh) or 10 nM 1,25D (blue, 1,25D).

**Table 1.** Patient Characteristics.

| Patient<sup>a</sup> | Age at RP<sup>b</sup> | Pathology | Fold change of organoid area with 1,25D | Significance of organoid area with 1,25D<sup>c</sup> | # of aliquots tested per patient | # of replicate wells per condition |
| --- | --- | --- | --- | --- | --- | --- |
| EA1 | 59 | benign | NA<sup>d</sup> | NA |  |  |
| EA2 | ? | benign | **1.41** | **0.0313** | 1 | 3 |
| EA3 | 57 | benign | **1.345** | **0.0011** | 2 | 6 |
| EA4 | 60 | benign | **1.708** | **<0.0001** | 3 | 9 |
| AA1 | 71 | benign | **1.359** | **<0.0001** | 4 | 12 |
| AA2 | 68 | benign | **1.22** | **0.0142** | 1 | 3 |
| AA3 | 72 | benign | NA | NA |  |  |
| AA4 | 58 | benign | **1.1119** | **0.0251** | 2 | 6 |
| AA5 | 60 | benign | **1.182** | **0.0019** | 2 | 6 |
| AA6 | 50 | benign | **1.47** | **0.0007** | 2 | 6 |
| AA8 | 58 | benign | **1.189** | **0.0536** | 1 | 3 |
| AA9 | ? | benign | NA | NA |  |  |
| Other1 | 58 | benign | 0.9680 | 0.4616 | 1 | 3 |

<sup>a</sup>EA = European American, AA = African American, Other = non-European or African descent
<sup>b</sup>Non-parametric one-tailed Mann Whitney test
<sup>c</sup>RP = radical prostatectomy
<sup>d</sup>NA = patient not used for organoid area assay

### Epithelial populations in primary prostate cell organoids by scRNAseq

Having established a 1,25D-induced phenotype of increased size across multiple patient-derived organoids (**Figure 1**), we next determined how 1,25D affected the epithelial populations within the organoids over time. Control and 1,25D-sufficent organoids from patient AA2 were collected for scRNAseq at two time points along the differentiation spectrum; day 8 and 14 (**Figure 2A,** left). AA2 was selected due to availability of cells from this patient, capacity for these cells to form acinar organoids, and response to 1,25D. Target 1,25D genes and pathways identified by scRNAseq in AA2 were validated in additional patients’ cells. Seurat version 3 was used to integrate datasets, align similar cells found in each sample, generate clusters, and perform differential expression analysis (Butler et al., 2018; Satija et al., 2015); the resulting UMAP of the integrated dataset and distribution of epithelial clusters found in each sample are shown (**Figure 2A,** middle & right). Cluster markers were determined by unbiased identification of genes that had uniquely high or low expression in each cluster (**Figure 2B, Supplemental Figure S2A**) and were input into Ingenuity Pathway Analysis (IPA) software to identify canonical pathways enriched in each population (Kramer et al., 2014) (**Supplemental Figure S2B**), as described below. To name the clusters, the expression of specific previously-identified epithelial markers was interrogated and visualized by dot plot (**Figure 2C**), which shows the percentage of cells in each cluster that express the marker of interest, and how high or low that expression is relative to the other cells in the dataset.

**Figure 2.**
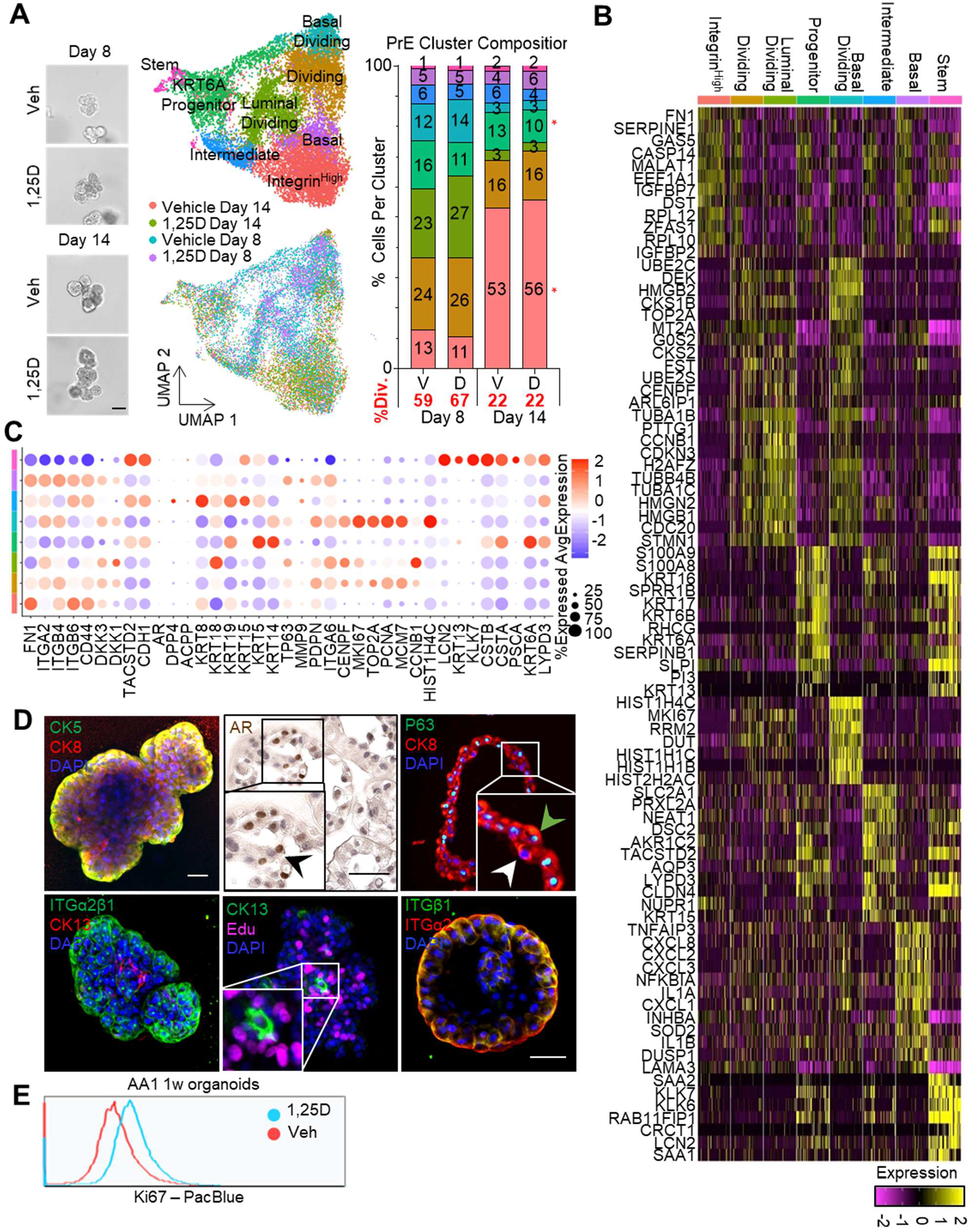
**Epithelial populations in primary prostate cell organoids by scRNAseq** (see also Figure S2). (A) Representative images of the AA2 organoids grown in vehicle (Veh) or 10 nM 1,25D (1,25D) collected at day 8 and day 14 for scRNAseq (left). Scale bar = 100 μm. UMAP of integrated scRNAseq data for vehicle and 1,25D samples at day 8 and day 14 (middle), showing clustering (upper) and distribution of samples (lower). Bar chart depicting the percentage of cells per cluster in each sample, vehicle (V) and 1,25D (D), at each time point (right), percentage of total dividing (%Div.) cells is shown in red. (B) Heat map of cluster markers for integrated scRNAseq dataset shown in 2A. Cluster markers were identified by Seurat as genes having uniquely high or low expression in each cluster, compared to all other cells in the dataset. (C) Dot plot showing expression of specific epithelial markers in each cell cluster. The size of dots represents the percentage of cells in each cluster that express the gene and the color represents relative expression of the cells in that cluster compared to all remaining cells. (D) Immunostaining of formalin-fixed, paraffin embedded organoids (upper middle and right) and whole-mounted organoids (remaining panels). Staining for luminal makers CK8 and AR, basal markers CK5 and P63, stem cell marker CK13, polarization markers ITGα2β1, ITGα2 and ITGβ1, and incorporation of Edu are shown. Scale bar = 50 μm. (E) Overlay of flow cytometry histogram for proliferation marker Ki67 in AA1 organoids grown until day 7 in vehicle (red, Veh) or 10 nM 1,25D (blue, 1,25D).

The monolayer cells from which the organoids originated consist primarily of transit-amplifiers with rare stem cells; our scRNAseq analysis revealed that with androgen stimulation and a 3D environment, the cells differentiate into eight clusters along the basal and luminal hierarchy. Resident, quiescent stem cells were identified by high expression of *KRT13* (Henry et al., 2018; Hu et al., 2017) (**Figure 2B & C**). This cluster had the most distinct expression profile with the highest number of markers (**Supplemental Figure S2A**). By IPA, the *KRT13* stem cell population had specific activation of the Wnt/β-catenin and VDR/RXR pathways, similar to what has been reported in gut and skin stem cells (Bikle, 2004; Peregrina et al., 2015) (**Supplemental Figure S2B,** red asterisks). Downstream progenitors were marked by modest *KRT13* expression but high expression of the progenitor marker *KRT6A* (Schmelz et al., 2005), along with expression of basal markers *KRT5/KRT14,* intermediate marker *KRT19* (Wang et al., 2001), and *TACSTD2* which codes for the stem cell marker Trop2 (Goldstein et al., 2008) (**Figure 2 B & C**). A luminal phenotype was distinguished by expression of *KRT8/KRT18,* and *DPP4* (CD26) (**Figure 2C**). Although, transcripts for *DPP4* were low and do not reflect the protein expression of CD26 observed by flow cytometry. Two basal populations were identified by *KRT5, KRT14, TP63* and *PDPN* (Henry et al., 2018) (**Figure 2C**). Dividing cells were recognized by cell division markers *CENPF, MKI67, TOP2A, PCNA, MCM7, CCNB1* and *HIST1H4C* (**Figure 2C**). The “Integrin^Hi^g^h^” population was identified based on integrin expression *(ITGB6, ITGB4,* **Figure 2C**), expression of integrin binding partner *FN1* (**Figure 2B**), and other adhesion junction proteins *(DST, CD44,* **Figure 2B & C**) (Ryan et al., 2012; Sneath and Mangham, 1998). Markers for the Integrin^High^ cluster were enriched for the HIPPO pathway, which regulates contact inhibition of polarized cell types (Genevet and Tapon, 2011) (**Supplemental Figure S2,** red asterisks). These cells may be starting to polarize through integrin interactions with the basement-membrane-protein ligands found in the Matrigel.

Immunostaining of organoids confirmed protein expression for major cluster markers. Basal proteins CK5 and p63 were expressed along the periphery of the organoid (**Figure 2D,** upper left & right green arrowhead). Although AR mRNA transcripts were low in the dataset, AR^+^ luminal cells were observed by IHC (**Figure 2D**, upper middle, black arrowhead). The luminal marker CK8/18 was highly expressed by most cells in the organoids, but there were p63^-^/CK8^+^ cells indicating a continuum of basal-intermediate-luminal cells in the organoids (**Figure 2D, upper left & right**, white and green arrows). This was also observed in the scRNAseq data where 100% of cells in every clusters express KRT8 (**Figure 2C**), but at different relative levels between the clusters. A CK13^+^ stem cell identified in an organoid was negative for EdU incorporation after briefly pulsing an organoid overnight with EdU prior to staining (**Figure 2D**, lower middle), supporting its quiescence. Integrin subunits α2, β1 and heterodimer integrin α2β1 were observed at the organoid–Matrigel interface where they are likely binding collagen and laminins found in the Matrigel to direct cell polarity (**Figure 2D**, lower left & right). CK13^+^ cells were negative for integrin α2/β1, demonstrating separation between the stem and Integrin^High^ clusters. In sum, these results confirm differentiation of a heterogeneous population of epithelial cells sharing markers identified by scRNAseq.

Once clusters were identified, population shifts over time and with vitamin D were explored using Fisher’s exact test (**Supplemental Figure S3A**, p value shown inset in the heat map). Early-stage day-8 organoids consisted primarily of dividing cell types, indicative of rapid cellular expansion in culture and significant compared to day 14 organoids (**Figure 2A,** red text, **Supplemental Figure S3A**). After differentiation at day 14, there was significant enrichment in the Integrin^High^ polarized cells compared to day 8. Culture in sufficient 1,25D increased the percentage of dividing cells at day 8 from 59% to 67% of the cell population (**Figure 2A**, red text, **Supplemental Figure S3A**), which was significant (**Supplemental Figure S3B**) and validated by flow cytometry for KI67 (**Figure 2E**). At day 14, population shifts with 1,25D were modest but significant (**Supplemental Figure S3C**), with a slight increase in the number of polarized and basal cells and a decrease in the number of progenitor and intermediate cells (**Figure 2A,** red asterisks). The data support that 1,25D promotes both growth and differentiation through enrichment of dividing cells at day 8 and Integrin^High^ cells at day 14.

### 1,25D modulates the Wnt pathway in organoids

Like most hormones, vitamin D is pleiotropic and regulates many pathways in prostate cells. To determine the action of 1,25D in organoids, differentially expressed genes (DEGs) after culture in 1,25D compared to vehicle in each cluster at each time point were explored (**Supplemental Figure S4A**). Of note, the stem cells had the fewest DEGs with 1,25D, indicating a stable transcriptional program. DEGs were input into IPA software to identify enriched Canonical Pathways (**Figure 3A and Supplemental Figure S4B**), Upstream Regulators (**Supplemental Figure S5**), and their Downstream Effects (**Supplemental Figure S6**) (Kramer et al., 2014). Although, the cluster shifts with vitamin D were subtle, they contain enrichment of key differentiation pathways, aligning with protective effects of the hormone, thus those were prioritized for investigation.

**Figure 3.**
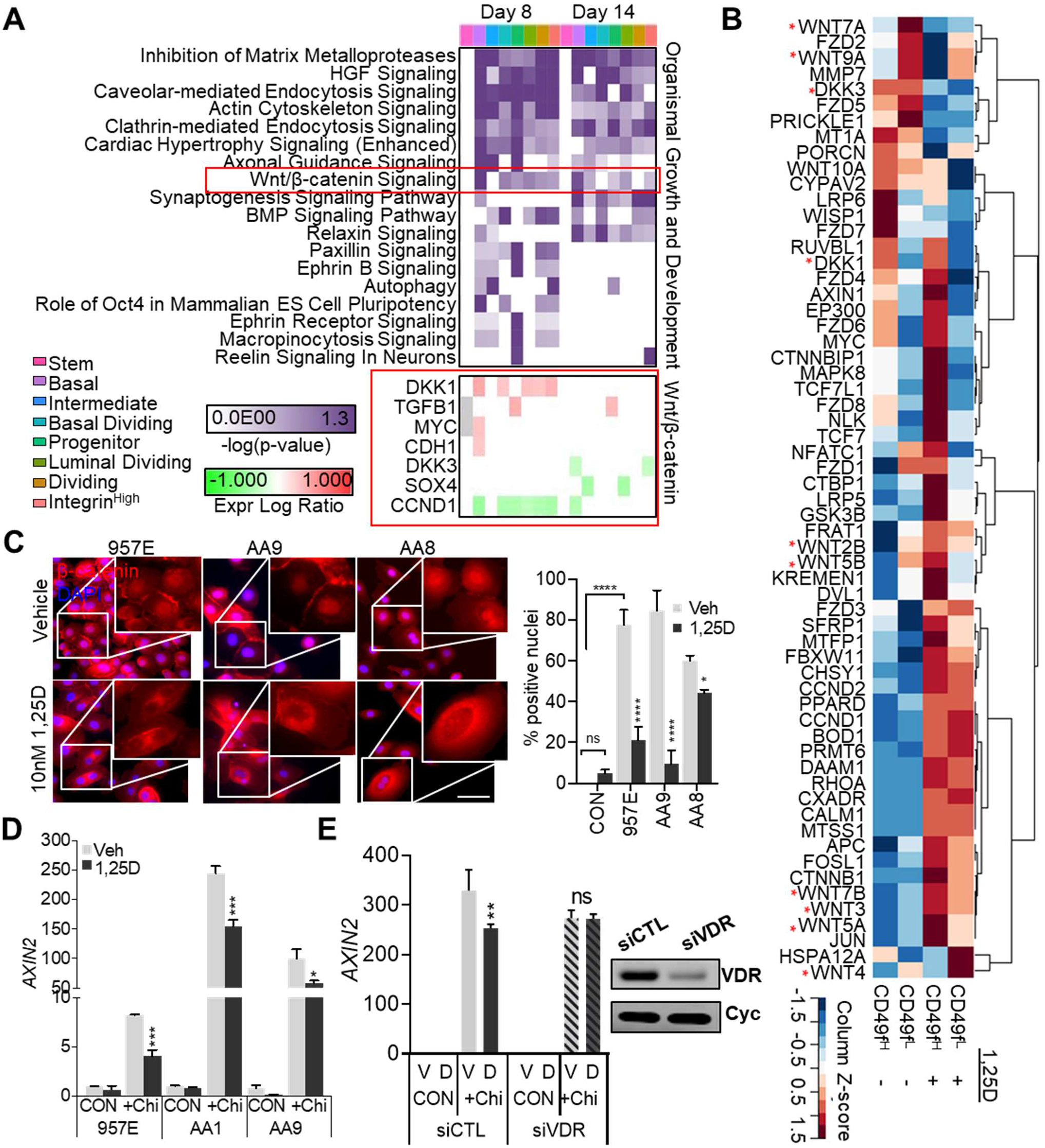
**1,25D modulates the Wnt pathway in organoids** (see also Supplemental Figures S3, S4, S5, S6). (A) Enriched pathways related to “Organismal Growth and Development” with 1,25D treatment in each cluster at each time point. Differentially expressed genes (DEGs) with 1,25D treatment per cluster were input into IPA canonical pathway analysis, pathways related to “Organismal Growth & Development” are shown. Scale represents significance, -log(p-value), for enrichment of each pathway with 1,25D. Dark purple is significant at p<0.05. Red box shows DEGs related to Wnt/β-catenin signaling with 1,25D treatment in each cluster. Scale represents the log ratio with 1,25D compared to control. (B) RT-qPCR array for Wnt pathway gene expression in flow-sorted day 17 AA3 organoids grown in vehicle or 10 nM 1,25D. Cells were sorted by CD49^High^ (CD49f^H^) or CD49^Low^ (CD49f^L^). Log2 of RQ values are shown, normalized to average of housekeeper genes. Hierarchical clustering was performed by Pearson correlation for row and by Spearman for column. (C) β-catenin protein localization in 1,25D-treated prostate epithelial cells. Monolayer PrE from two patients (middle & right panel) and benign 957E (left panel) cells grown for 48 or 96 h, respectively, with vehicle or 10 nM 1,25D. Cells were treated with 9 μM Chiron for 5 h, followed by immunostaining for β-catenin (red) and counterstaining with DAPI (blue). Scale bar = 50 μm. Quantification of the percentage of positive nuclei is shown (right), error bars represent the standard deviation of two fields of view. The average of the -Chiron controls (CON, treated with DMSO) is shown to illustrate significant nuclear B-catenin induction upon Chiron treatment (AA8 CON was not collected). P value represents outcome of paired 2-way ANOVA with uncorrected Fisher’s comparison by row for vehicle vs. 1,25D, and for CON vs. 957E Chiron Veh. (ns = not significant, *p<0.05, **** *p* < 0.0001). (D) RT-qPCR of *AXIN2* expression in monolayer PrE cells and benign 957E cells grown and treated as in (C). Relative quantitation is normalized to *HPRT1.* Error bars represent standard deviation of technical RT-qPCR pipetting replicates. Cells from three sources serve as biological replicates. P value represents outcome of paired 2-way ANOVA with uncorrected Fisher’s comparison by row for vehicle vs. 1,25D (*p < 0.05; ***p < 0.001). (E) RT-qPCR of *AXIN2* expression in monolayer AA1 PrE cells transfected with siCTL or siVDR RNA and grown as described in (C)(left). Relative quantitation is normalized to *HPRT1.* Error bars represent standard deviation of technical RT-qPCR pipetting replicates. P value represents outcome of paired 2-way ANOVA with uncorrected Fisher’s comparison by row for vehicle vs. 1,25D. Western for VDR in siCTL and siVDR samples (right) with housekeeper cyclophilin-B (cyclo), full western blot included in **Supplemental Figure S7**.

Significantly enriched canonical pathways in all clusters included VDR/RXR activation (**Supplemental Figure S4B**), driven by the highly 1,25D-regulated genes *CYP24A1* and
*IGFBP3* (**Supplemental Figure S4B,** upper red box) (Martin and Pattison, 2000; Zierold et al., 1995). Examination of pathways related to “Organismal Growth & Development” (**Figure 3A**) showed enrichment of the BMP and Wnt pathways in 1,25D sufficient organoids, known regulators of prostate development and differentiation (Toivanen and Shen, 2017). DEGs within the Wnt/β-catenin signaling pathway included upregulation of *DKK1* at day 8 and downregulation of *DKK3* at day 14 (**Figure 3A**, lower red box). Stem cells showed the fewest number of DEGs with 1,25D culture (**Supplemental Figure S4A**) and this cluster showed no enrichment for pathways related to “Organismal Growth & Development.” Vitamin D status did not alter the stem population viability as shown by three passages of self-renewal assays in four patients (**Supplemental Figure S4C**). Together, this indicates that 1,25D mainly regulates lineage-committed cells on the path to differentiation and does not alter the stable transcriptional program in the stem cells.

Upstream Regulator analysis predicted active transcription factors which may have influenced DEG expression. Activation of VitaminD3-VDR-RXR and cell-type-specific regulation by CTNNB1 were among the top regulators (**Supplemental Figure S5,** red asterisks), confirming vitamin D function and further supporting Wnt/β-catenin involvement. To understand the net effect of the Pathway and Regulator analysis, downstream Disease & Function analysis was performed (**Supplemental Figure S6**). Consistent with the hypothesis, there was enrichment for “Differentiation of Epithelial Tissue” at both time points with sufficient 1,25D (**Supplemental Figure S6,** red box). The Wnt pathway was selected for additional investigation in the primary prostate cells because previous studies in the RWPE1 prostate cell line, gut, skin and heart have shown 1,25D inhibition of canonical Wnt signaling (Aguilera et al., 2007; Hlaing et al., 2014; Kovalenko et al., 2010; Larriba et al., 2011; Muralidhar et al., 2019).

Wnt regulation by 1,25D was validated with a targeted RT-qPCR array (**Figure 3B**). Prior to RNA isolation, organoids cultured with 10 nM 1,25D or vehicle were sorted by high-or low-CD49f (Guo et al., 2012; Yamamoto et al., 2012) in an attempt to preserve differences in transcription between stem vs lineage-committed cells or basal vs luminal cells. The expression of multiple canonical and non-canonical Wnts were altered by 1,25D, in some cases in a CD49f- cell-type-specific manner, including: *WNT7A, WNT9A, WNT2B, WNT5B, WNT7B, WNT3, WNT5A,* and *WNT4.* Notably, *DKK3* was downregulated with 1,25D treatment in both high (CD49f^H^) and low cells (CD49f^L^) (**Figure 3B**, red asterisks).

To determine overall directionality of Wnt regulation by 1,25D, β-catenin translocation and *AXIN2* induction were assessed. The GSK3β-inhibitor, Chiron, was used for β-catenin stabilization and Wnt pathway induction. PrE cells from two patients (AA8, AA9) and the benign 957E cell line grown with 1,25D had decreased nuclear β-catenin after Chiron treatment compared to vehicle controls (**Figure 3C**). Consistent with reduced β-catenin activity, 1,25D abrogated Chiron-induced activation of the Wnt response gene *AXIN2* (**Figure 3D**). Knockdown of VDR in AA1 monolayer cells (**Figure 3E, Supplemental Figure S7A-C**), blocked 1,25D’s ability to inhibit *AXIN2* after Chiron treatment (**Figure 3E**). Overall, these data support an inhibitory effect of 1,25D on the canonical Wnt pathway and implicate direct regulation via VDR.

### 1,25D inhibits Dickkopf family member 3 (DKK3)

*DKK3* emerged as a 1,25D target in both the scRNAseq dataset and the RT-qPCR array, so its expression was profiled in a panel of prostate cell lines, monolayer PrE cells, and organoids (**Figure 4A**). All the cell lines, both immortalized benign (RWPE1 and 957E) and PCa (LAPC4 and PC3) had very low or undetectable *DKK3* expression. DKK3 is also known as “Reduced Expression in Immortalized Cells” (REIC) (Hsieh et al., 2004), thus our findings are consistent with previous reports of limited expression in cell lines. Across PrE cells grown as monolayers or organoids, 1,25D inhibited *DKK3* expression (**Figure 4A**).

**Figure 4.**
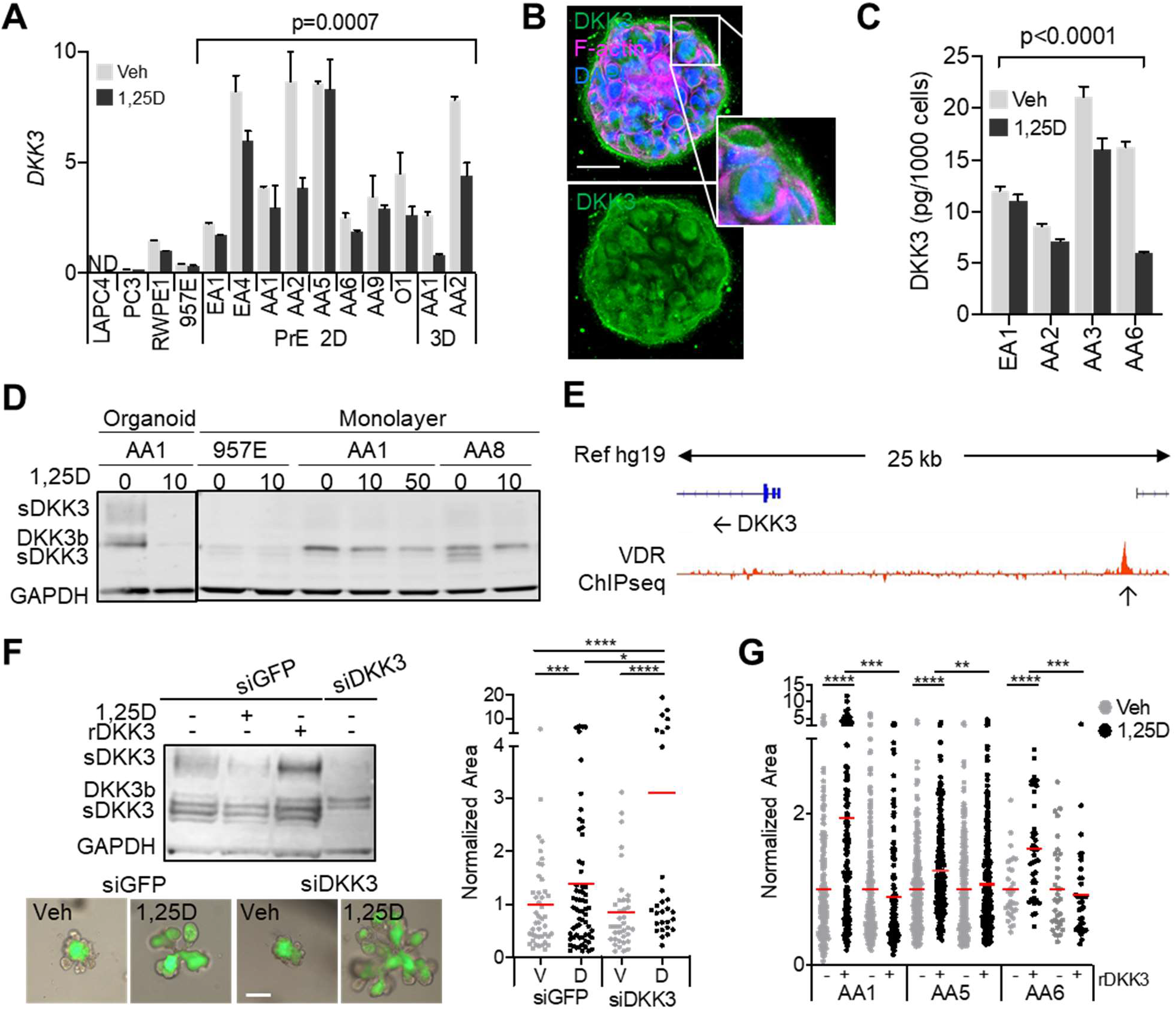
1,25D inhibits Dickkopf family member 3 (DKK3). (A) RT-qPCR of *DKK3* expression in prostate cell lines (LAPC4, PC3, RWPE1, 957E), monolayer prostate epithelial cells (PrE 2D), and organoid PrE cells (3D) grown in vehicle or 10 nM 1,25D. Monolayer cells were treated for 24–72 h, and organoids were treated for 2–3.5 weeks. Relative quantitation is normalized to *HPRT1;* error bars represent standard deviation of technical RT-qPCR pipetting replicates. Cells from multiple sources represent biological replicates. P value represents interaction outcome of 2-way ANOVA comparison for vehicle vs 1,25D in PrE cells. (B) Immunostaining for DKK3 (green) in a whole-mounted day-17 AA3 organoid counterstained with DAPI (blue) and phalloidin/F-actin (pink). Scale bar = 50 μm. (C) ELISA quantification of secreted DKK3 in media collected from monolayer PrE cells grown in vehicle or 10 nM 1,25D for 72 h, normalized to the number of cells. Error bars represent the standard error of triplicate ELISA technical samples. Cells from multiple sources present biological replicates. P value represents interaction outcome of 2-way ANOVA comparison of vehicle vs 1,25D. (D) Western blot of DKK3 expression in cell lines and monolayer and organoid PrE cells grown in vehicle or 10 nM 1,25D (sDKK3 = secreted DKK3 variants; DKK3b = intracellular DKK3 variant). (E) VDR-ChIP sequencing peak near DKK3 in PrE cells treated with 1,25D for 3 hours, IGV track showing 1,25D-treated minus vehicle-treated coverage (data from NCBI GEO accession number GSE124576). (F) Western blot of monolayer AA1 prostate epithelial (PrE) cells transduced with control virus (siGFP) or siDKK3 virus and grown in vehicle, 10 nM 1,25D, or 50 ng/mL rDKK3 (left). Representative images of siGFP-or siDKK3-treated AA1 PrE organoids grown in vehicle or 10 nM 1,25D for 2 weeks (middle), with quantification of relative area, normalized to vehicle (right) (*p < 0.1; **p < 0.05, ***p < 0.01; ****p < 0.001. Scale bar = 200 μm). (G) Relative area of PrE organoids grown in vehicle or 10 nM 1,25D combined with 50 ng/mL rDKK3 treatment for two weeks, normalized to vehicle (*p < 0.1, **p < 0.05; ***p < 0.01; ****p < 0.001).

Immunofluorescence for DKK3 revealed a vesicular and perinuclear staining pattern (**Figure 4B,** white box), similar to other secreted members of the Dickkopf family (Glinka et al., 1998; Inoue et al., 2017). ELISA showed that culture in 1,25D reduced DKK3 secretion in media (**Figure 4C**), and western blots of cell lysates showed reduced intracellular protein levels with 1,25D treatment (**Figure 4D**). Western blot of DKK3 detected several sized bands, agreeing with preceding reports describing many variants: a heavy secreted-form (sDKK3), a lighter intracellular variant (DKK3b), and a 50-kDa secreted form (sDKK3) (Abarzua et al., 2005; Hsieh et al., 2004; Kawano et al., 2006; Leonard et al., 2017; Zenzmaier et al., 2008; Zhang K. et al., 2010). The 957E cell line had no detectable expression of sDKK3 and faint expression of DKK3b, uniform with the low RNA expression observed. PrE cells grown as monolayers and organoids showed reductions in all variants of DKK3 with sufficient 1,25D compared to controls (**Figure 4D**). Analysis of our previously published VDR-ChIP-seq dataset (Baumann et al., 2019) in PrE cells showed a peak 20 kb upstream from DKK3 after 1,25D treatment (**Figure 4E**), supporting potential genomic regulation by VDR.

To emulate 1,25D actions, PrE cells were transduced with lentivirus containing an siDKK3 sequence and GFP tag (**Figure 4F, left**) and grown as organoids in the presence of vehicle or 10 nM 1,25D (**Figure 4F, lower left**). Knockdown alone did not recapitulate the effect of 1,25D treatment, but combination of siDKK3 with 1,25D significantly enhanced the effect of vitamin D on organoid area (**Figure 4F,** right). To rescue 1,25D inhibition of DKK3, exogenous recombinant DKK3 (rDKK3) was added to culture. Treatment with rDKK3 enhanced all DKK3 variants within cell lysates (**Figure 4F, left**). PrE organoids from three patients were grown in media supplemented with vehicle, 1,25D, or 1,25D in combination with 50 ng/mL rDKK3 for 2 weeks. rDKK3 did not affect organoid area under vehicle conditions, but blocked the increase in area seen in 1,25D-treated organoids (**Figure 4G**). In sum, these data support that inhibition of DKK3 by 1,25D serves to promote organoid growth.

### 1,25D regulates DKK3 in lineage-committed cells

The function of DKK3 in the Wnt pathway is not well understood, but it has been shown to inhibit proliferation in prostate and breast cancer cell lines (Leonard et al., 2017). Expression of *DKK3* and *KRT13* was mutually exclusive in the scRNAseq dataset, suggesting that *DKK3* is expressed only in lineage-committed cells (**Figure 5A,** lower). To observe lineage-committed cells and separate them from progenitor and stem cells, the scRNAseq data was subset into three groups based on *KRT13* expression (**Figure 5B**, left). *DKK3* was expressed at both time points and most visibly inhibited by 1,25D at day 14 (**Figure 5B**, right), indicating stronger inhibition with increased length of exposure. Immunofluorescence for CK13 and DKK3 in organoids showed no colocalization (**Figure 5C**). To observe expression across organoid differentiation trajectories, day-8 and day-14 vehicle scRNAseq datasets were plotted together in pseudotime using Monocle 3 (**Figure 5D**). *KRT13* was expressed at the beginning of pseudotime, with cells from day 8 and day 14 samples found at this stage, showing the resident stem cells in organoid culture (Hu et al., 2017)(**Figure 5C**, upper right & lower left). Day-8 cells clustered halfway through the differentiation trajectory where *MKI67* expression is high (**Figure 5D,** upper right & lower middle), and day-14 cells clustered at the end (**Figure 5D**, upper right). A fork was seen in day-14 cells, possibly where basal and luminal lineages start to diverge. *DKK3* expression increased across pseudotime, consistent with expression in lineage-committed cells and indicating its upregulation in senescence (Untergasser et al., 2002) (**Figure 5C**, bottom right).

**Figure 5.**
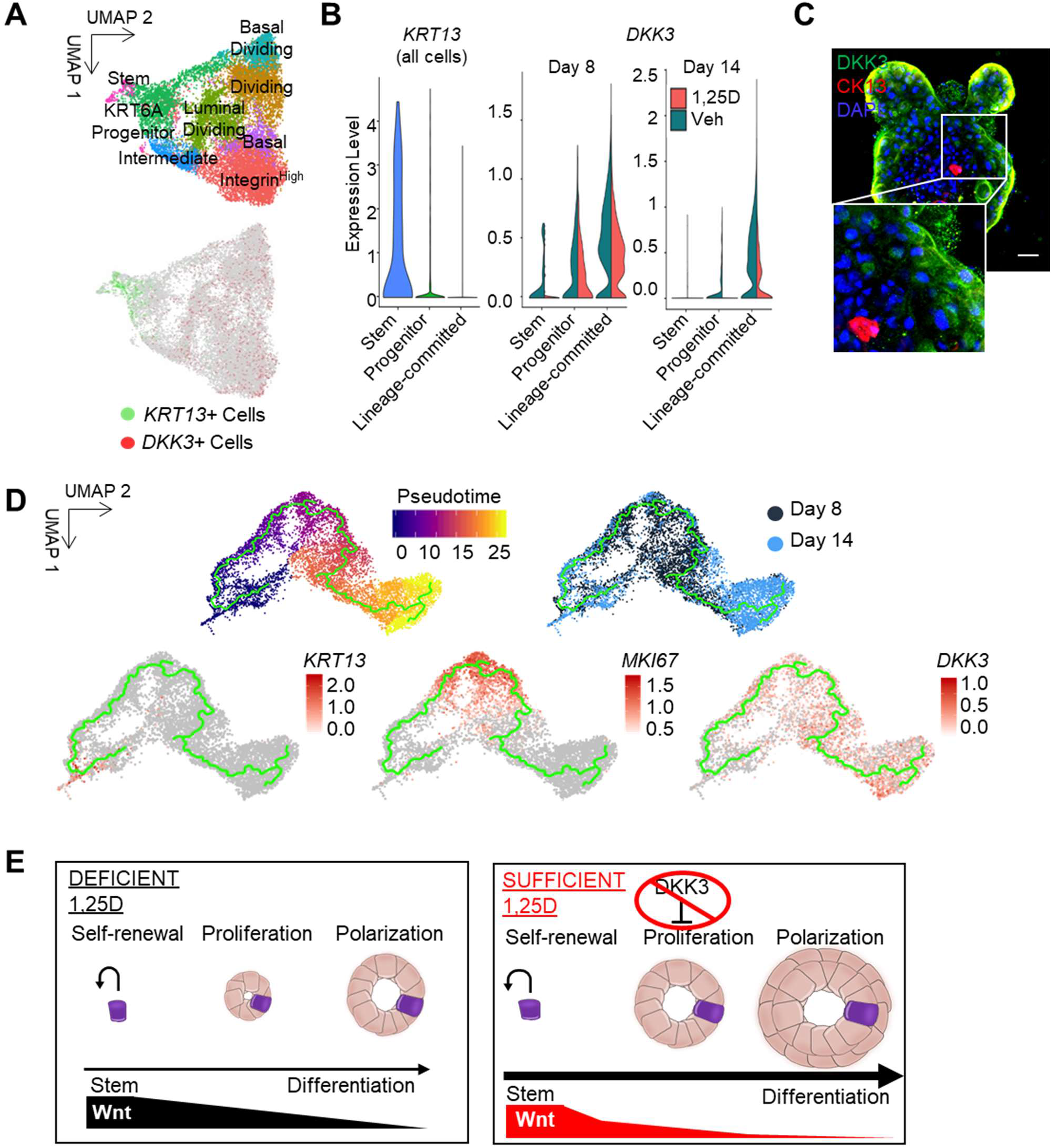
1,25D inhibits DKK3 in lineage-committed cells. (A) UMAP of integrated scRNAseq dataset shown in Figure 2A (upper), blended UMAP of integrated scRNAseq data showing expression of *KRT13* (green) and *DKK3* (red) (lower). (B) Violin plot of integrated dataset shown in (A), subset by *KRT13* expression into lineage-committed, progenitor or stem populations (left). Violin plots of integrated dataset shown in 5A, separated by time point, showing *DKK3* expression (right) in vitamin D (pink) and vehicle (teal) samples. (C) DKK3 (green) and CK13 (red) expression in whole-mounted AA3 organoid counter-stained with DAPI (blue). Scale bar = 50 μm. (D) Pseudotime of integrated scRNAseq data for day-8 and day-14 vehicle samples (upper left). Sample distribution across pseudotime for day-8 (dark blue) and day-14 (light blue) cells (upper right). *KRT13* (lower left), *MKI67* (lower middle), and *DKK3* (lower right) expression across pseudotime. (E) Diagram summarizing the effects of 1,25D on prostate epithelial organoids. Under vehicle conditions “DEFICIENT 1,25D”, a self-renewing stem cell with high Wnt activity undergoes asymmetric division to produce a progenitor cell that will rapidly expand. As differentiation occurs, downstream cells will have reduced Wnt activity. Under sufficient conditions, “SUFFICIENT 1.25D”, proliferation is enhanced through inhibition of DKK3, resulting in a larger organoid. Wnt signaling is also inhibited in lineage-committed cells through β-catenin sequestration, to promote differentiation away from a stem cell phenotype.

## DISCUSSION

The prostate is a hormonally regulated gland that requires steroids for development, and dysregulation of hormones occurs during carcinogenesis and late-stage PCa. Vitamin D is a steroid hormone that promotes differentiation in many cell types, yet its role had not been studied in the differentiation of benign human prostate epithelium. Using primary epithelial organoids from multiple patients, we found that continuous culture in physiologically relevant concentrations of 1,25D promoted differentiation and increased organoid area compared to vitamin D-deficient conditions (**Figures 1 & 2**). The dominant mechanisms were inhibition of the canonical Wnt pathway (**Figure 3**) and reduction of Wnt pathway member DKK3 (**Figure 4 & 5**), to promote epithelial growth of lineage-committed cells (**Figure 5E**).

The Wnt pathway is known to be highly active in prostate stem cells compared to differentiated cells (Blum et al., 2009). In kidney organoids and prostate organogenesis, Wnt activity is critical early-on to promote progenitor outgrowth but then decreases to allow for differentiation (Prins and Putz, 2008; Simons et al., 2012; Takasato et al., 2016). Similarly, in snake venom gland organoids, Wnt agonists must be removed to allow for differentiation and secretory function (Post et al., 2020). Consistently, we observed that lineage-committed cells had lower Wnt activity and selective Wnt inhibition by 1,25D was observed in the lineage-committed cells, but not in *KRT13+* stem cells, resulting in enhanced epithelial differentiation in the organoids (**Figure 5E**). This effect of vitamin D complements the reports of Wnt pathway regulation in other epithelial models. Kovalenko et al reported that 1,25D inhibited Wnt and promoted genes that “induced during differentiation” from Gene Set Analysis (GSA) in the benign prostate RWPE1 cell line (Kovalenko et al., 2010). In colon cancer cell lines and mouse colon tissue, 1,25D inhibited the Wnt pathway by decreased nuclear β-catenin and increased cellular differentiation (Aguilera et al., 2007; Groschel et al., 2016; Larriba et al., 2013). Similar results of β-catenin by vitamin D metabolites were observed in Kaposi’s sarcoma (Tapia et al., 2020) and renal cell carcinoma (Xu et al., 2020).

In PCa, Wnt signaling has been shown to promote resistance to androgen deprivation therapy (Yokoyama et al., 2014). PCa stem cells show increased nuclear β-catenin and TCF/LEF activity (Zhang et al., 2017) and Lef-1 identifies an androgen-insensitive population of basal progenitors in mouse prostate maturation (Wu et al., 2011). As PCa progresses, *APC* and *CTNNB1* become mutated in 22% of castration resistant PCa to drive Wnt pathway activation (Murillo-Garzon and Kypta, 2017). In PCa cell lines, disruption of e-cadherin enhanced Wnt signaling and increased tumor growth (Davies et al., 2000) and WNT5A has been shown to promote cancer cell invasion (Wang et al., 2020). Our results indicate that vitamin D sufficiency could provide negative pressure on the Wnt pathway to complement other therapies and improve PCa patient outcome.

DKK3 emerged as potently and consistently inhibited by 1,25D in the prostate organoids. Our findings suggest an anti-proliferative role for DKK3 in prostate organoids and this adds to literature that is inconsistent about the downstream target of this protein in prostate cells. In general, the Dickkopf family members are inhibitors of Wnt signaling, such as DKK1 (Glinka et al., 1998; Kruithof-De Julio et al., 2013). However, DKK3 is the most structurally divergent member of the Dickkopf family (Krupnik et al., 1999) and has varied effects on Wnt, ranging from no effect (Krupnik et al., 1999; Pinho Christof Niehrs, 2007; Romero et al., 2013) to promoting (Ferrari et al., 2019; Nakamura and Hackam, 2010; Yin et al., 2018) or inhibiting Wnt (Bhattacharyya et al., 2017; Leonard et al., 2017; Liu et al., 2019; Sharma Das et al., 2013). DKK3 also functions as both a positive and negative regulator of TGFβ signaling, depending on the model (Al Shareef et al., 2018; Busceti et al., 2017; Kardooni et al., 2018; Li et al., 2017; Pinho Christof Niehrs, 2007; Romero et al., 2013; Wang Z et al., 2015), but DKK3 did not impact TGFβ in this study (data not shown). Despite conflicting reports of DKK3’s signaling targets, it has consistently been shown to restrain cell proliferation (Kawano et al., 2006; Leonard et al., 2017). It acts as a cell cycle inhibitor whose expression is reduced in immortalized cells (Kawano et al., 2006), upregulated in senescent PrE cells at passage 10 compared to passage 3 (Untergasser et al., 2002) and upregulated during aging (Yin et al., 2018). In developing mouse prostate, addition of exogenous DKK3 blunted proliferation, preventing luminal differentiation, Nkx3.1 expression, and epithelial bud formation (Kruithof-De Julio et al., 2013). Our findings that 1,25D reduction of DKK3 resulted in increased organoid size is similar to the DKK3 deficient mice, which are viable with normal prostate glands, but have increased Ki-67+ proliferating cells (Romero et al., 2013). We observed that DKK3 was not expressed by the stem cells in organoids, similar to patient tissue (Henry et al., 2018), indicating that DKK3 acts downstream to regulate cell growth (**Figure 5E**). Notably, DKK3 is also highly expressed in the stroma of the prostate, which indicates a multifaceted role for this protein that can be explored in the future using co-culture models (Al Shareef et al., 2018; Henry et al., 2018; Zenzmaier et al., 2013). The increase in organoid size induced by 1,25D may seem contradictory to the traditional anti-proliferative role of Vitamin D metabolites (Feldman et al., 2014; Murphy et al., 2017; Wagner et al., 2013). However, the organoid model was optimized to create a permissive environment for expansion and differentiation of stem/progenitor cells and is supplemented with androgen to drive cell division.

Access to primary cells from multiple, ancestrally-diverse patients and the use of scRNAseq are major strengths of this study, yet they are not without limitations. Patient-derived cells have inherent inter-patient heterogeneity and epithelial content in the tissue collected, as a result the aliquots available differs between patients. To address this, we used as many patients as possible for each endpoint and supplemented with cell lines if necessary. ScRNAseq is a powerful tool to analyze the transcriptomes of thousands of cells from heterogenous samples, but it only captures a subset of the transcripts in each cell (∼30-32% of transcripts for the 10x Chromium Single cell v3 kit). Thus, the lack of a transcript does not mean that protein is absent, for example *DPP4* and *AR* were low in the scRNAseq data, but CD26 was detected by flow cytometry, nuclear AR was detected by IHC and CK8^+^/p63^-^ cells were observed (**Figure 1E, Figure 2D**). Given that organoids grow from the AR-negative stem cells found within monolayer PrE culture, the presence of basal cells and AR+ luminal cells confirms some differentiation. As well, because the organoids were completely epithelial without other cell types, the data lack extremely different clusters. However, our clusters were transcriptionally distinct and similar to those observed in scRNAseq data from the epithelial clustering of whole patient prostate tissue (Henry et al., 2018). Despite these limitations, the approach successfully identified vitamin D-regulated genes and pathways that were validated by other assays.

In summary, we report two complementary mechanisms by which vitamin D deficiency disrupts prostate epithelial differentiation—inhibition of canonical Wnt signaling and the protein DKK3. This is the first report, to our knowledge, of the hormone vitamin D regulating DKK3 expression in the prostate. These findings are particularly impactful for patients who are frequently deficient in vitamin D, such as African Americans and adults over the age of 70, who have significantly high rates of PCa (Jacques et al., 1997; Murphy et al., 2014).

## ACKNOWLEDGEM ENTS

We thank UIC Biorepository (Alex Susma), and urologists (Drs. Michael Abern, Daniel Moreira, and Simone Crivallero) for facilitation of tissue acquisition for patient-derived cultures. We thank UIC Urology patients for donating their tissue—without their participation, none of this research would be possible. We thank Dr. Ke Ma with UIC Fluorescence Imaging Core for assistance with confocal. We thank Dr. Matt McDougall of the Merrill laboratory for his counsel with the Wnt pathway. We thank Dr. Alvaro Hernandez, Chris Wright, Jenny Zadeh, and Jessica Holmes and the UIUC DNA services team for sequencing support and the CellRanger pipeline. We thank Magdalena Rogozinska and Dr. Gayatry Mohapatra for their help with TapeStation quantification. We thank UIC Pathology Department members Joseph Marsili, Morgan Zenner, Jason Garcia, Larischa DeWet, and Lenny Hong for their thoughts and troubleshooting. We thank Junbin Huang for his participation on the project through ResearcHStart. This work was funded, in part, by the Department of Defense Prostate Cancer Research Program Health Disparities Idea Award PC121923 (Nonn) and the UIC Center for Clinical and Translation Science Pre-Doctoral Education for Clinical and Translational Scientists (PECTS) Program (McCray). The content is solely the responsibility of the authors and does not necessarily represent the official views of the National Institutes of Health or Department of Defense.

## AUTHOR CONTRIBUTIONS

Conceptualization, T.M and L.N.; Formal analysis, T.M.; Investigation, T.M., J.P., B.B., M.J.S; Resources, L.N., K.V.N.; Writing, T.M., B.B.; Visualization, T.M.; Supervision, L.N.; Funding acquisition, L.N.

## METHODS

### LEAD CONTACT AND MATERIALS AVAILABILITY

Further information and requests for resources and reagents should be directed to and will be fulfilled by the Lead Contact, Larisa Nonn. This study did not generate new unique reagents.

### EXPERIMENTAL MODEL AND SUBJECT DETAILS

#### Primary Cell Culture

Primary prostate cells were generated from fresh human male radical prostatectomy samples as previously described (Giangreco and Nonn, 2013; Nonn et al., 2006). Briefly, tissue from benign regions of the peripheral zone were collected according to an IRB approved protocol of UIC Cancer Center Tissue Bank Core Facility (protocol # 2004-0679). A board-certified pathologist confirmed the tissue portion as benign from adjacent H&E section. Tissue was digested in collagenase/trypsin, and single cells were plated in Prostate Cell Growth Media (Lonza, Basel, Switzerland) to enrich for epithelial cells. At ∼70% confluence, cells were frozen into multiple aliquots and cryopreserved for experiments. Epithelial purity was confirmed by RT-qPCR for expression of epithelial markers *KRT5, KRT8, KRT18,* and *TP63* (data not shown). Patient information is listed on **Table 1**. All cells were cultured at 37°C with 5% CO_2_. Due to differences in patients’ cells growth capacity and availability of cells, different patients were used for different endpoints.

#### Cell Lines

RWPE1 cells were purchased from ATCC in 2014, cryopreserved in low-passaged aliquots, reauthenticated in 2016, and used only at <20 passages. LAPC4 were generously donated from Dr. Charles Sawyers and grown in RPMI 1640 (Gibco, MD) with 10% fetal bovine serum on polylysine-coated flasks. 957E/hTERT cells were generously donated from Dr. Don Vander Griend. RWPE1 and 957E cells were grown in uncoated cell culture dishes in Keratinocyte Serum Free Media (Corning, NY). PC3 were purchased from ATCC, re-authenticated in 2013, and grown in RPMI 1640 (Gibco, MD) with 10% fetal bovine serum. All cells were cultured at 37°C with 5% CO_2_.

#### Organoid Culture

Organoids were grown as previously described by our group (McCray et al., 2019b; Richards et al., 2019). Briefly, epithelial cells were thawed and grown to passage 2 on collagen-coated dishes to ∼70% confluency. Epithelial cells were plated singly and sparsely (100–1,000 cells per well, depending on patient-specific growth ability) in 100 μL of 10%–50% growth factor-reduced Matrigel (Corning, NY). To prevent cells from adhering to the bottom and growing as a monolayer, cells were plated over a 50% Matrigel base layer in a 96-well plate or low-attachment 96-well plate without a base layer. Matrigel was diluted with organoid growth media containing Keratinocyte Serum Free Media (Gibco, MD), bovine pituitary extract, and epidermal growth factor, supplemented with 5% charcoal-stripped fetal bovine serum and 10 nM dihydrotestosterone (DHT). After Matrigel solidified, 100 μL of organoid growth media was added to each well and was refreshed every 1–3 days. Media was supplemented with vehicle or 1,25D, and cells as indicated in figure legends. Given the patient heterogeneity in growth capacity, cells were collected at subtly different time points.

### METHOD DETAILS

#### Cell Treatments

Monolayer and organoid primary cells and monolayer cell lines were maintained in media as described above. Cells were treated with vehicle or 10 nM or 50 nM 1,25D over the course of culture, as indicated in figure legends. For Wnt induction, cells were treated with vehicle or 9 μM Chiron for 5 h before collection. For rDKK3 (1118-DK-050, R & D Systems, MN), cells were exposed to 50 ng/mL over the course of culture, as indicated in figure legends.

#### Brightfield Image Acquisition, Processing, and Analysis

Images of organoid cultures were acquired and analyzed as previously described by our group (McCray et al., 2019b; Richards et al., 2019). Briefly, images were captured at 4x magnification using the Evos FL Auto 2 Imaging System (Thermo Fisher Scientific, MA), and up to 50 z-planes were captured per quadrant of a 96-well plate. Images of each quadrant were stitched together at each z-position, and z-stacks were compressed to an enhanced depth-of-field image using Celleste Image Analysis software to visualize each organoid in the well within a single image (Thermo Fisher Scientific, MA). Images were also collected at 10x and 20x of individual organoids for better resolution for figures.

##### Organoid morphology

Solid, translucent and acinar organoids were counted per well for n ≥ 3 wells of each condition for 10 patients, and the percentage of each morphology was calculated. Percentage average and standard error of the mean was determined for the 7 patients that were capable of forming all three types of morphologies.

##### Organoid number and area

Organoid count and area metrics were generated by manual identification of each organoid using Celleste Image Analysis software. At least three wells per patient were analyzed for technical replicates, each patient had 1-4 biological replicates performed by thawing vials for repeat experiments from each patient, and each patient serves as a biological replicate for the phenotype. Area was measured at roughly day 14, +/− 4 days. Area was normalized to vehicle control, and a non-parametric, one-sided unpaired, Mann Whitney t-test was used to compare between treatments, p<0.1 was considered significant.

#### Self-renewal assay

Replating self-renewal assay was performed to assess stem cell sphere formation ability. Epithelial cells were plated singly and sparsely (100–1,000 cells per well, depending on patientspecific growth ability) in 100 μL per well of 10% growth factor-reduced Matrigel (Corning, NY) on a low-attachment 96-well plate without a base layer. Matrigel was diluted with organoid growth media containing Keratinocyte Serum Free Media (Gibco, MD), bovine pituitary extract, and epidermal growth factor, supplemented with 5% charcoal-stripped fetal bovine serum and 10 nM dihydrotestosterone (DHT). After Matrigel solidified, 100 μL of organoid growth media was added to each well and was refreshed every 1-3 days. Media was supplemented with vehicle or 1,25D, and cells were grown for 5-7 days in culture. After spheres formed, they were dissociated into single cells using TryPLE and replated in the same conditions for another passage, up to passage 3. Cells were imaged at the day of plating to determine the number of cells seeded and imaged at day 5-7 to determine the number of spheres formed. The % of spheres formed was determined by dividing the number of spheres by the number of cells seeded.

#### Immunostaining

Immunostaining was performed as previously described by our group (McCray et al., 2019b). Organoids were collected from Matrigel by Dispase (Stemcell Technologies, MA), resuspended in HistoGel (Thermo Fisher Scientific, MA), fixed in 4% paraformaldehyde, and paraffin-embedded. Whole-mounted organoids were transferred by pipette to a chamber slide, fixed in 4% paraformaldehyde, and permeabilized. GFP signal in transduced cells was quenched with 50 mM NH4Cl if 488-channel was used for immunostaining. Sections (5 μm) of embedded organoids or whole-mounted organoids were incubated overnight at 4°C with rabbit monoclonal anti-p63a antibody (D2K8X, Cell Signaling Technology, MA), polyclonal guinea pig anti-cytokeratin 8/18 (03-GP11, American Research Products Inc, MA), monoclonal rabbit anti-cytokeratin 13 (ab92551, Abcam, UK), monoclonal rabbit anti-androgen receptor (D6F11, Cell Signaling Technology, MA), monoclonal rabbit anti-□-catenin (8480, Cell Signaling Technology, MA), monoclonal mouse anti-integrin α2/β1 (ab20483, Abcam, UK), polyclonal rabbit anti-keratin 5 (905501, BioLegend, CA), or monoclonal rabbit anti-DKK3 (ab186409, Abcam, UK). Samples were incubated with Alexa Fluor conjugated secondaries for 1 h at room temp or overnight at 4°C. Samples were counterstained with Alexa Fluor 647-phalloidin and DAPI, when appropriate. For whole-mount staining of cytokeratin 13 and DKK3, incubation with unconjugated monoclonal rabbit anti-DKK3 (ab186409, Abcam, UK) and secondary antibody was performed, followed by incubation with conjugated monoclonal rabbit anti-cytokeratin 13 (EPR3671 conjugated to Alexa Fluor 647, ab198585, Abcam, UK) and counterstaining. Edu staining was performed as previously described (McCray et al., 2019a).

#### Immunostaining Quantification

Positive and negative nuclei for □-catenin staining were manually identified from two fields of view images for each condition. Number of nuclei was identified from DAPI and positive nuclei were identified by localization of red □-catenin for each cell in the field of view.

#### Lentivirus Transduction

Cells were transduced by centrifugation (Xin et al., 2003) with DKK3 siRNA/shRNAi Lentivirus (iV006166) or scrambled siRNA GFP Lentivirus (LVP015-G) (Applied Biological Materials Inc., CA) and selected by passaging at least once into media containing 1–3 μg/mL puromycin until all cells were GFP+ before plating into Matrigel.

#### SiRNA Transfection

PrE AA1 cells were transiently transfected as previously described (Blajszczak and Nonn, 2019). Briefly, cells were plated on a 60 mm dish and transfected with a pool of three 19-25 siRNA sequences against VDR (sc-106692) or control siRNAs (sc-37007) (Santa Cruz Biotech, TX) using DharmaFECT Duo Transfection Reagent (T-2005-01) (Horizon Discovery, UK). Cells were plated and given 24 hours to attach and divide. At 24 h, siRNAs were added and incubated for 48 h to allow siRNA incorporation and inhibition of VDR translation. After 48 h, a second siRNA delivery was performed along with 1,25D treatments. Cells were collected after 48 h with 1,25D, and 96 h with siRNAs. Prior to RNA collection, Chiron was added for 5 h for the *AXIN2* assay, as described in the cell treatments section. Protein was also collected to validate knockdown.

#### Cell Sorting

Day-14 organoids from AA1 were dissociated with Dispase and TrypLE Express Enzyme (Gibco, MA) to generate single-cell suspensions and were fixed with ethanol. Cells were incubated with Alex Fluor 647-conjugated monoclonal rat anti-CD49f (BioLegend, CA), PE-conjugated monoclonal mouse anti-CD26 (BioLegend, CA), and PacificBlue-conjugated monoclonal mouse anti-Ki67 (BioLegend, CA) antibodies and were sorted with LSR Fortessa (BD Biosciences, CA). FlowJo software (FlowJo LLC, OR) was used to create scatterplot overlays of samples at each time point.

For live cell sorting for RT-qPCR array, day-17 organoids were dissociated as described above and incubated with Alex Fluor 647-conjugated monoclonal rat anti-CD49f (BioLegend, CA) and FITC-conjugated monoclonal mouse anti-CD26 (BioLegend, CA) antibodies. Cells were live-sorted with the MoFlo sorter (Beckman-Coulter, CA) directly into TRIzol (Thermo Fisher Scientific, MA) for RNA isolation.

#### PCR Profiling Arrays and RT-qPCR gene expression

For PCR profiling array, organoid RNA was isolated using TRIzol extraction. RNA quantity and quality were determined by NanoDrop Spectrophotometer (Thermo Fisher Scientific, MA). cDNA was generated with Qiagen RT^2^ First Strand Kit (Qiagen, Germany), and gene expression was assessed using RT^2^ Profiler PCR Array Human WNT Signaling Pathway Plus (PAHS-043YC-2, Qiagen Germany). Arrays were run on QuantStudio6 (Thermo Fisher Scientific, MA) and normalized independently using five reference genes according to the manufacturer’s protocol. The △△CT method was used for comparative analysis (Livak and Schmittgen, 2001).

For individual RT-qPCR, RNA was isolated with TRIzol and reverse-transcribed using a High Capacity cDNA Reverse Transcription Kit (Thermo Fisher Scientific, MA), qPCR was performed on the QuantStudio6 machine with SYBR green. Expression of *VDR* was quantified using forward primer 5’-GACCTGTGGCAACCAAGACT-3’ and reverse primer 5’-GAACTTGATGAGGGGCTCAA-3’. Expression of *CYP24A1* was quantified using forward primer 5’-CATTTTAGCAGTCAGCTCCCG-3’ and reverse primer 5’-GGCAACAGTTCTGGGTGAAT-3’. Expression of *AXIN2* was quantified using forward primer 5’-AGTGTGAGGTCCACGGAAAC-3’ and reverse primer 5’-ACAGGATCGCTCCTCTTGAA-3’. Expression of *DKK3* was quantified using forward primer 5’-TCACATCTGTGGGAGACGAA-3’ and reverse primer 5’-CTGGCAGGTGTACTGGAAGC-3’. Gene expression was normalized to the reference gene *HPRT1* with forward primer 5’-TGCTGACCTGCTGGATTACA-3’ and reverse primer 5’-CTGCATTGTTTTGCCAGTGT-3’.

#### Western Blot

Organoids were collected from Matrigel, lysed in cell lysis buffer (9803, Cell Signaling, MA) with protease/phosphatase inhibitor (5872, Cell Signaling, MA), and sonicated. Bradford assay was used to detect protein concentration. Protein was denatured before separation on 1.5 mm 4%–12% NuPAGE Bis-Acrylamide gels with NuPAGE MOPS SDS running buffer and NuPage antioxidant (Thermo Fisher Scientific, MA). Gels were primed for 10 min at 100 V and run for 1 h at 125 V. Proteins were transferred from gels to 42-μm-pore Millipore PVDF membranes (Sigma-Aldrich, MO) for 1 h at 30 V. Protein membranes were blocked using Odyssey Blocking Buffer (LI-COR Biosciences, NE). Primary antibodies were diluted in Odyssey Blocking Buffer and incubated with blots overnight at 4°C. Primary antibodies used include monoclonal mouse anti-cyclophilin-B (CL3901, Abcam, UK), monoclonal rabbit anti-VDR (D2K6W, Cell Signaling, MA), monoclonal rabbit anti-DKK3 (ab186409, Abcam, UK), and monoclonal mouse anti-GAPDH (MAB374, Millipore Sigma, MA). Secondary antibodies IRDye 800CW anti-mouse IgG (926-32210, LI-COR Biosciences, NE) and IRDye 680RD anti-rabbit IgG (926-68071, LI-COR Biosciences, NE) were diluted in Odyssey Blocking Buffer and incubated with blots for 1 h at room temperature. Blots were imaged using the LI-COR Odyssey Imaging system.

#### DKK3 ELISA

Primary epithelial cells were grown as monolayers and treated with vehicle or 10 nM 1,25D for 72 h, with treatments refreshed every 24 h. At 48 h, cells were washed, and media depleted in bovine pituitary extract was added to prevent detecting exogenous bovine DKK3. Media was collected at 72 h, and cells were counted with a Cellometer Automated Cell Counter (Nexcelom, MA) for normalization. Protein was quantified using Human Dkk-3 DuoSet Solid Phase Sandwich ELISA (DY1118, R&D Systems, MN), per manufacturer’s instructions. Plates were read using Synergy HTX Multi-mode Read (BioTek, VT).

#### scRNAseq

Organoids were collected by Dispase and TrypLE as described above, and scRNAseq was performed as previously described (McCray et al., 2019a). Briefly, cell number and viability were determined using the Cellometer Automated Cell Counter (Nexcelom, MA) by Trypan Blue exclusion. Samples were >85% viable cells, and ∼5,000 cells per sample were captured with the 10x Chromium chip (10x Genomics, CA). Libraries were prepared per manufacturer’s instructions using the 3’ Transcript Capture and Single Cell Library Prep v3 chemistry (10x Genomics, CA). Libraries were quantified and titrated by MiSeq (Illumina, CA) into an even pool and sequenced across one 2Ø150nt lane of the NovaSeq 6000 (Illumina, CA) at a depth of ∼50,000 reads per cell. Library titration and sequencing was performed by University of Illinois at Urbana-Champaign (UIUC) DNA services. scRNAseq samples were processed and aligned to Ensembl genome GRCh38 using the Cell Ranger 3.2.1 pipeline (10x Genomics, CA) by UIUC DNA Services.

#### Seurat Integration and Clustering Analysis

The CellRanger output was loaded into Seurat 3.1.0 for clustering, following integration and differential gene expression workflows (Butler et al., 2018; Satija et al., 2015). Poor-quality cells with high mitochondrial gene expression and unusually high or low reads were subset out of the dataset to remove dying cells and doublets. Individual samples were normalized, cell cycle and mitochondrial features were scored, and data was scaled. Next, samples were integrated to find similar cells across samples, and cell cycle and mitochondrial features were regressed. Highly variable features were used for principal component analysis and reduction for UMAP clustering. We reduced 30 principal components at a resolution of 0.4 to obtain a UMAP plot with a modularity of 0.8249. Clusters were assigned epithelial identities based on expression of known epithelial markers, as previously described (McCray et al., 2019a). Cluster markers were determined by DEGs in each cluster compared to all remaining cells, determined by Seurat default nonparametric Wilcoxon rank sum test (Butler et al., 2018). Similarly, DEGs with 1,25D treatment compared to vehicle controls were determined for each cluster at both time points in the same fashion. Genes with adjusted *p* < 0.05 after Bonferroni correction were considered significant. Log2 odds ratio (color) and corresponding p-value (number inset) were determined from Fisher’s exact test comparing each cluster to all other clusters between conditions.

#### Monocle Pseudotime Analysis

CellRanger outputs for day-8 and day-14 vehicle data were loaded into Monocle 3 and combined, and pseudotime trajectories were constructed following the Trapnell lab’s workflow (Cao et al., 2019; Qiu et al., 2017; Trapnell et al., 2014). Briefly, data were loaded and normalized, samples were combined, batch effects were removed, cells with high mitochondrial gene expression were removed, and trajectories were constructed. The beginning node was selected based off of *KRT13* expression.

#### Ingenuity Pathway Analysis (IPA)

Lists of cluster markers and DEGs with 1,25D treatment that were generated in Seurat were analyzed with IPA (Qiagen, https://www.qiagenbioinformatics.com/products/ingenuitypathway-analysis). Gene lists were input into the core analysis function to determine canonical pathways, Diseases & Functions, and Upstream Regulators. Only cluster markers and DEG genes with adjusted p < 0.05 from Seurat’s default non-parametric Wilcoxon rank sum test were used. Once each core analysis was finished, a comparative analysis was performed for each pathway, function or Upstream Regulator across each cluster at each time point.

#### VDR ChIP Sequencing

ChIP sequencing for VDR-bound DNA near the *DKK3* promoter was observed in a previously published dataset of 1,25D-treated PrE cells from our lab (Baumann et al., 2019), deposited at NCBI GEO accession number GSE124576.

### QUANTIFICATION AND STATISTICAL ANALYSIS

Statistical analyses were performed with GraphPad Prism version 8 (GraphPad Software Inc., CA), Microsoft Excel (Microsoft Windows, WA), Seurat R Package (Butler et al., 2018), and IPA software (Qiagen Bioinformatics, DK); details can be found in figure legends. A non-parametric, one-sided unpaired, Mann Whitney U-test was used to compare organoid area. For RT-qPCR and ELISA analyses a 2-way ANOVA was used to compare conditions, standard deviation of replicates or standard error of mean is depicted by error bars as noted in the figure legends. Differential expression analysis was performed with Seurat, using a non-parametric Wilcoxon rank sum test, and adjusted Bonferroni corrected *p* < 0.05 was considered significant. P < 0.05 was used as a significant for IPA enriched pathway analysis.

### DATA AND CODE AVAILABILITY

ScRNAseq data has been submitted to NCBI Gene Expression Omnibus and can be found under accession number GSE142489. ChIP sequencing data can be found under accession number GSE124576.

