## Supplementary material for "Vitamin D Sufficiency Enhances Differentiation of Patient-Derived Prostate Epithelial Organoids": SuppLegends&Figs

### SUPPLEMENTAL INFORMATION TITLES AND LEGENDS

**Supplemental Figure S1, related to Figure 1: PrE express VDR and organoids respond to 1,25D**

(A) RT-qPCR of basal *VDR* expression in patient-derived prostate epithelial cells. Relative quantitation shown normalized to *HPRT1* (EA, European American; AA, African American; ancestries were patient self-declared). Error bars represent standard deviation of technical RT-qPCR pipetting replicates.

(B) RT-qPCR of *CYP24A1* expression in AA2 organoids cultured in vehicle, 10 nM or 50 nM 1,25D. Relative quantitation (RQ) shown normalized to *HPRT1.* Error bars represent standard deviation of RT-qPCR technical pipetting replicates, they are included but are small and difficult to visualize on scale of graph.

**Supplemental Figure S2 (related to Figure 2): Pathway enrichment analysis for cluster markers.**

(A) The number of markers per cluster. Cluster markers were identified by Seurat as genes having uniquely high or low expression in each cluster, compared to all other cells in the dataset. Markers that are highly expressed compared to all remaining cells are shown red, markers that are lesser expressed are blue.
(B) Cluster markers were input into IPA canonical pathway analysis. Significantly enriched pathways related to “Nuclear Hormone Signaling” and “Organismal Growth & Development” are shown. Dark purple is significant at $p<0.05$ for enrichment of each pathway (left) and activation z-score for each pathway (right).

**Supplemental Figure 3 (related to figure 2):** Log2 odds ratio (color) and corresponding p-value (number inset) from Fisher's exact test comparing each cluster to all other clusters between conditions. For example, the log2 odds ratio of 4 for Luminal Dividing cells (top right) means that that cells have a 16x higher odds of being part of this cluster at day 8 compared to day 14.

(A) Comparison of day 8 and day 14 vehicle clusters.
(B) Comparison of vehicle and 1,25D clusters at day 8.
(C) Comparison of vehicle and 1,25D clusters at day 14.

**Supplemental Figure S4 (related to Figure 3): Pathway enrichment analysis for Nuclear Hormone Signaling Pathways in differentially expressed genes with 1,25D treatment per cluster, per time point.**

(A) The number upregulated (red) and downregulated (blue) differentially expressed genes (DEGs) with 1,25D compared to vehicle in each cluster at day 8 (left) and day 14 (right).
(B) For each cluster and time point, DEGs with 1,25D were input into IPA canonical pathway analysis. Enrichment for pathways related to "Nuclear Hormone Signaling" are shown. Scale represents –log(p-value) for enrichment of each pathway with 1,25D treatment (left) and expression log ratio with 1,25D treatment compared to vehicle (right). Red box shows differentially expressed genes related to VDR/RXR activation. Dark purple is significant at $p<0.05$.
(C) % of organoids formed at passage 3 from self-renewal assay for spheres grown in vehicle or 1,25D and split every 5-7 days. % of organoids formed was calculated by dividing the # spheres at day 5 by the # cells at plating for passage 3. Each dot represents the outcome for a patient, 4 separate patients were tested. NS = not significant by Mann-Whitney t-Test for vehicle vs. 1,25D.

**Supplemental Figure S5 (related to Figure 3): Upstream regulator analysis for differentially expressed genes with 1,25D treatment per cluster per time point.** Top 70 significant regulators from IPA upstream regulator analysis of differentially expressed genes with 1,25D treatment per cluster. Scale represents predicated activation z-score.

**Supplemental Figure S6 (related to Figure 3): Diseases & Functions Enrichment analysis for differentially expressed genes with 1,25D treatment per cluster per time point.** Differentially expressed genes with 1,25D per cluster per time point were input into IPA Disease and Functions analysis. Significantly enriched Diseases & Functions related to “Molecular and Cellular Function, Physiological System Development and Function” are shown. Red box shows differentially expressed genes related to “Differentiation of Epithelial Tissue”. Scale represents activation z-score for enrichment (left) and expression log ratio with 1,25D compared to control (right).

**Supplemental Figure S7 (related to Figure 3):** siCTL and siVDR knockdown PrE cells

(A) RT-qPCR of *VDR* expression in AA2 and AA1 monolayer PrE cells transduced with CTL or VDR siRNAs twice across 96 hours. Relative quantitation (RQ) shown normalized to *HPRT1* and scaled to siCTL veh sample per patient. Error bars represent standard deviation of RT-qPCR technical pipetting replicates. P value represents the outcome of a 2-way ANOVA for siCTL vs siVDR, **p < 0.01.

(B) RT-qPCR of *CYP24A1* expression in AA2 and AA1 monolayer PrE cells transduced with CTL or VDR siRNAs twice across 96 hours. At 48 hours, cells were treated with vehicle or 10 nM 1,25D and cultured for an 48 h prior to RNA collection. Relative quantitation (RQ) shown normalized to *HPRT1* and scaled to the vehicle sample of the siCTL virus. Error bars represent standard deviation of RT-qPCR technical pipetting replicates. P value represents the outcome of a 2-way ANOVA for vehicle vs. 1,25D, ****p < 0.0001.

A

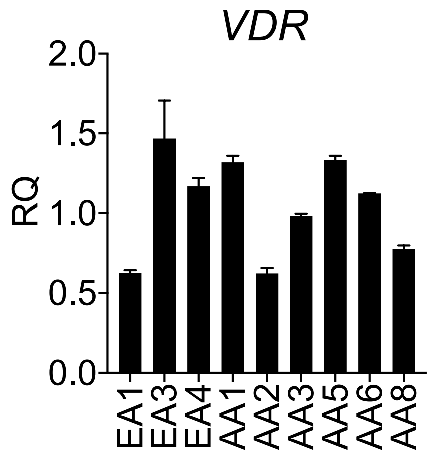


B

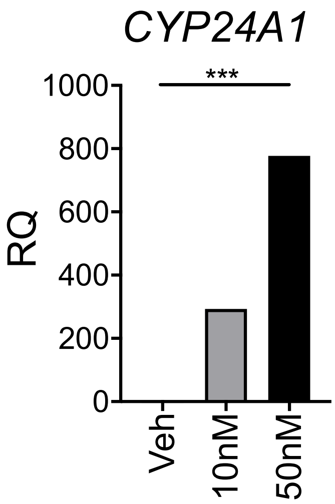


**Supplemental Figure 1, related to Figure 1**

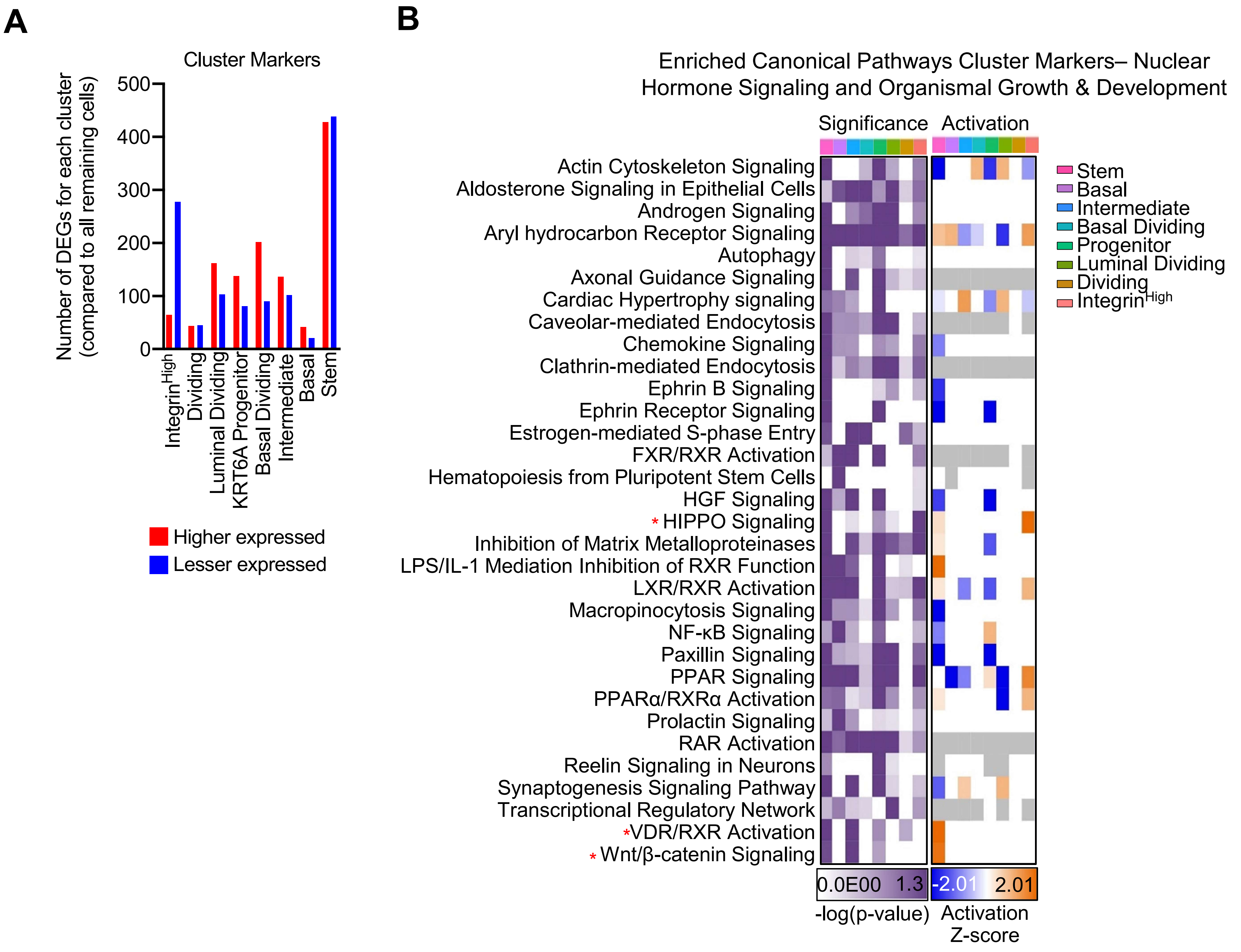


**Supplemental Figure 2, related to Figure 2.**

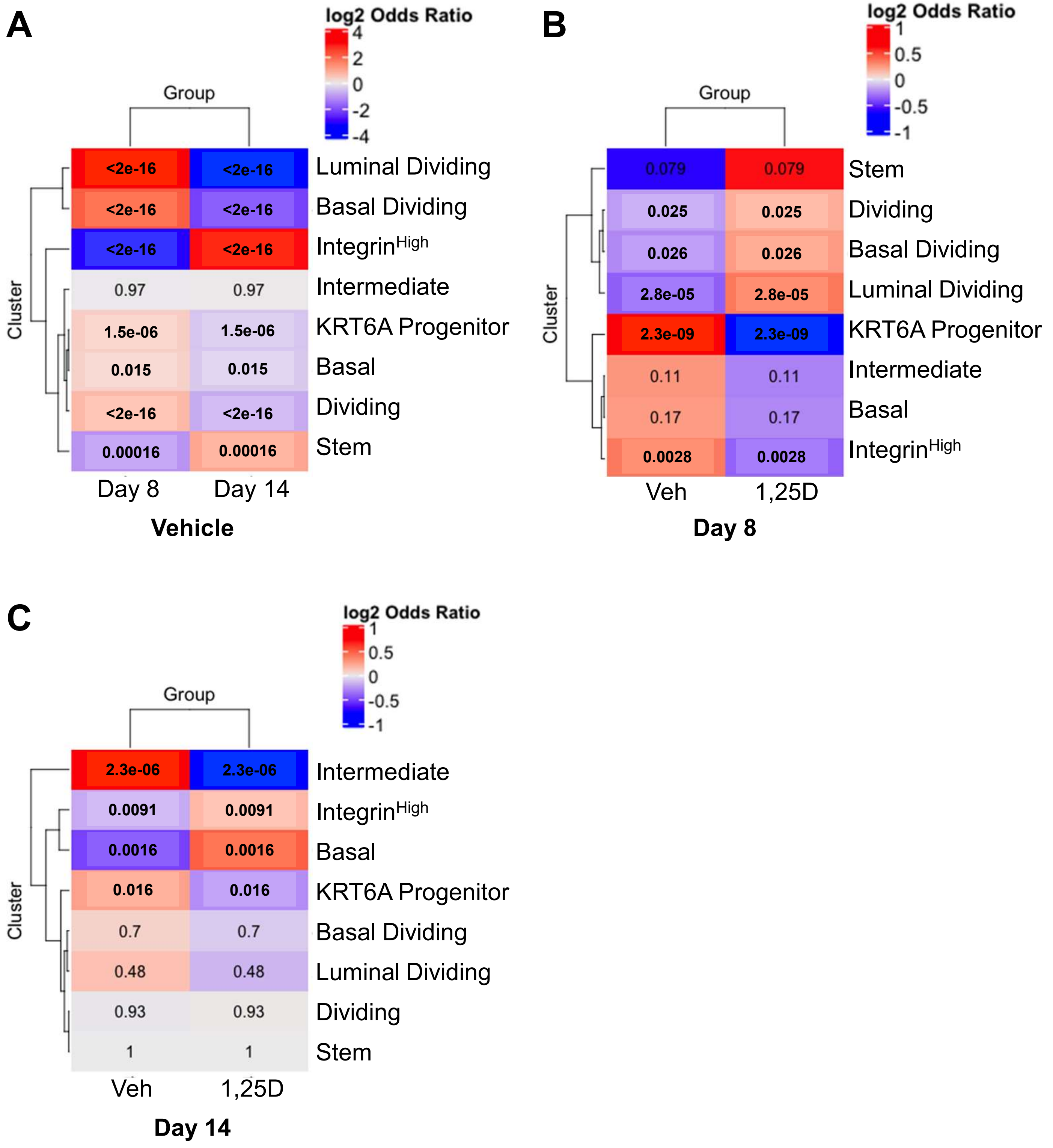


**Supplemental Figure 3 (related to figure 2)**

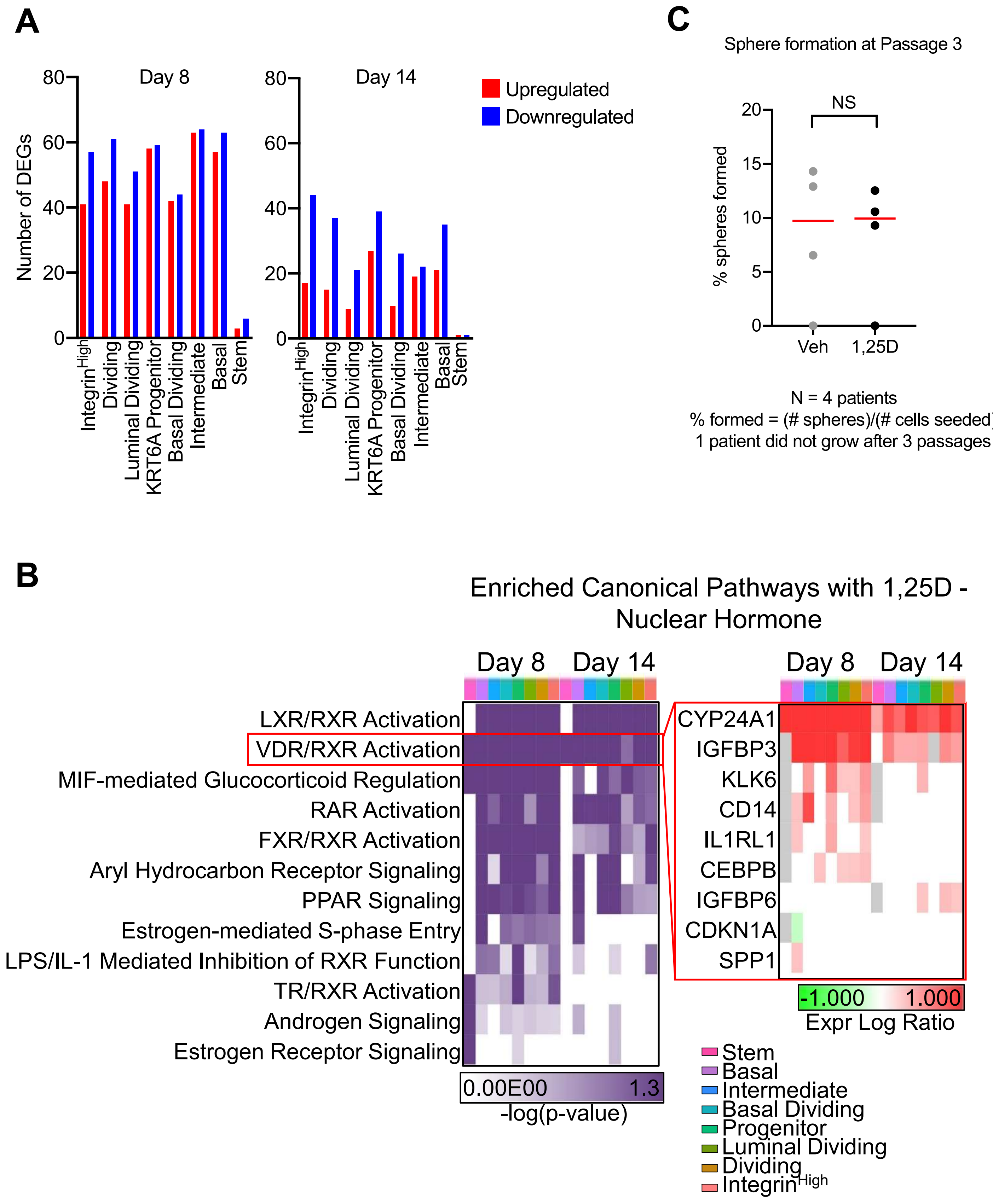


**Supplemental Figure 4 (related to figure 3)**

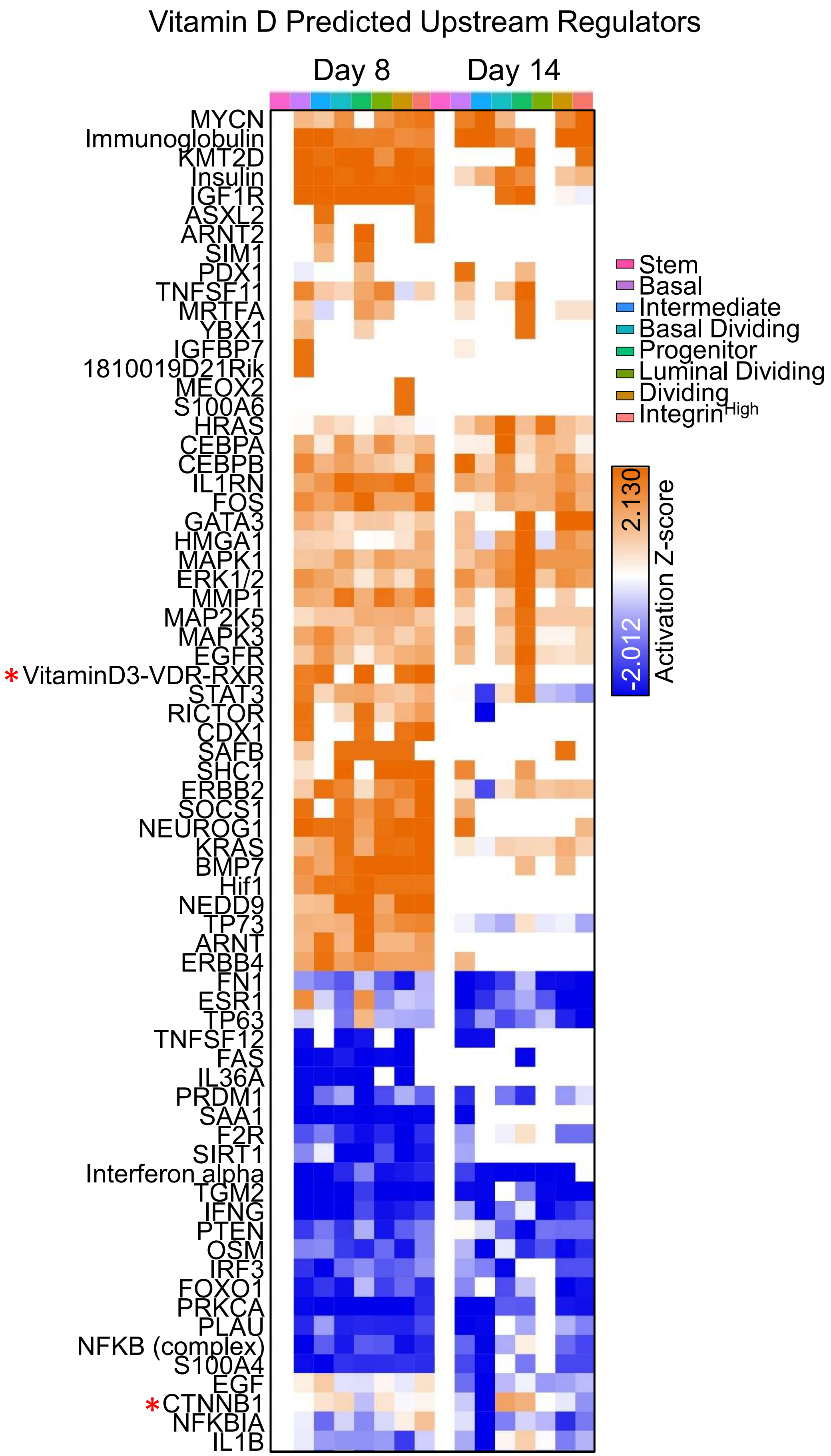


**Supplemental Figure 5 (related to figure 3)**

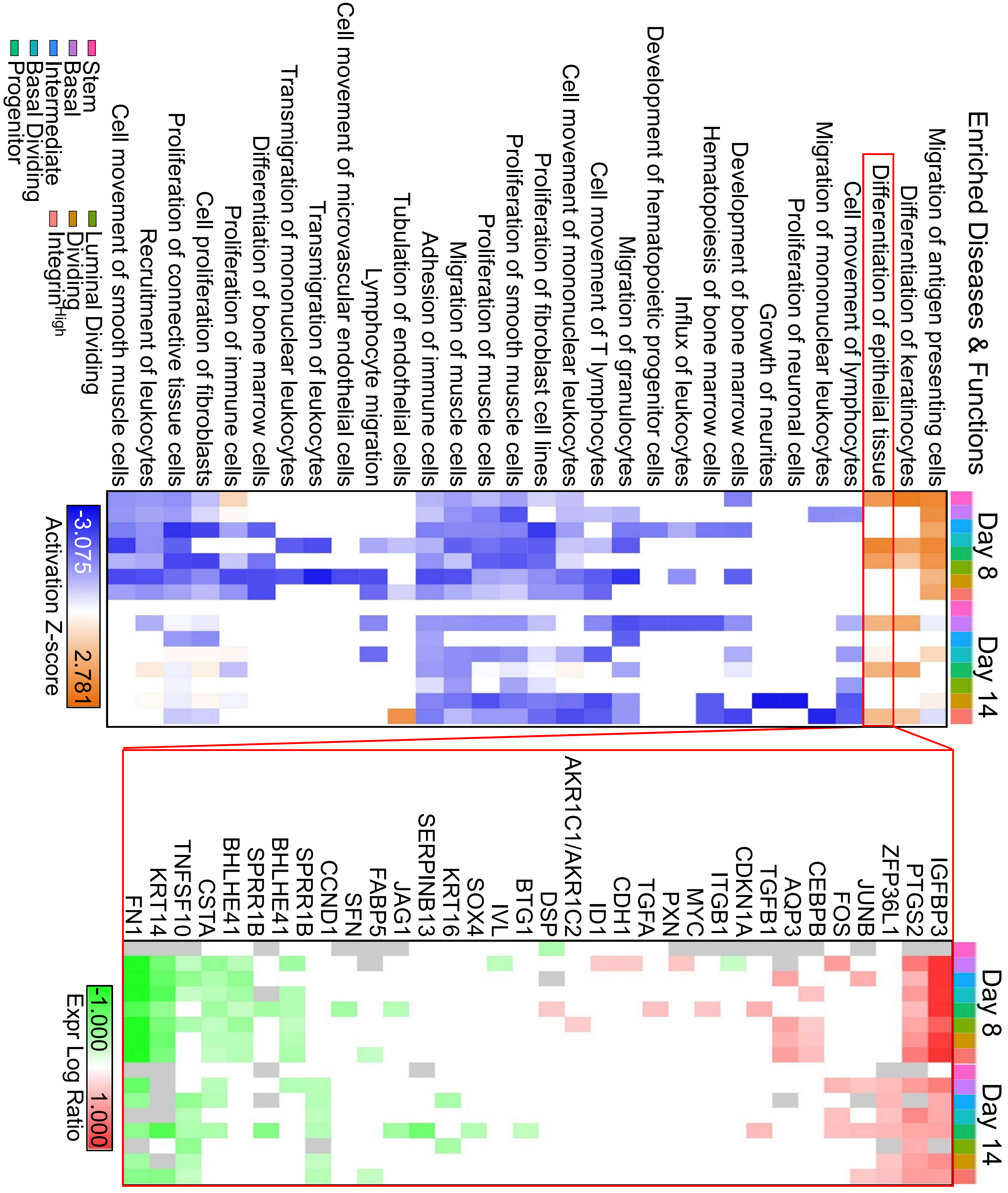


**Supplemental Figure 6 (related to figure 3)**

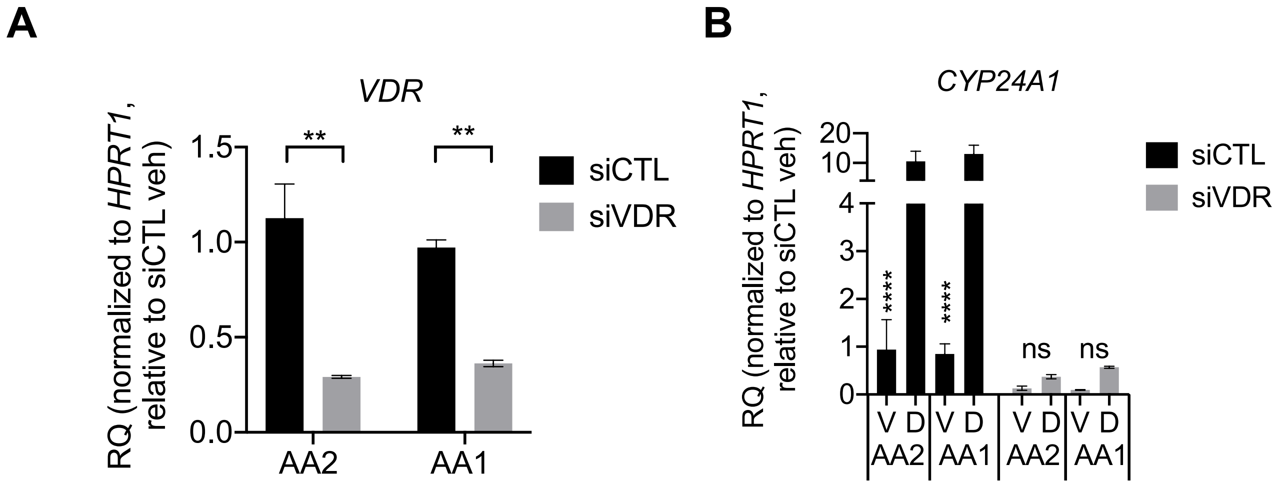


**Supplemental Figure 7 (related to figure 3)**
